# Staphylococcal aconitase expression during iron deficiency is controlled by an sRNA-driven feedforward loop and moonlighting activity

**DOI:** 10.1101/2024.05.23.595409

**Authors:** Maxime Barrault, Svetlana Chabelskaya, Rodrigo H. Coronel-Tellez, Claire Toffano-Nioche, Eric Jacquet, Philippe Bouloc

## Abstract

Pathogenic bacteria employ complex systems to cope with metal ion shortage conditions and propagate in the host. IsrR is a regulatory RNA (sRNA) whose activity is decisive for optimum *S. aureus* fitness upon iron starvation and for full virulence. IsrR down-regulates several genes encoding iron-containing enzymes to spare iron for essential processes. Here we report that IsrR regulates the tricarboxylic acid (TCA) cycle by controlling aconitase (CitB), an iron-sulfur cluster-containing enzyme, and its transcriptional regulator, CcpE. This IsrR-dependent dual-regulatory mechanism provides an RNA-driven feedforward loop, underscoring the tight control required to prevent aconitase expression. Beyond its canonical enzymatic role, aconitase becomes an RNA-binding protein with regulatory activity in iron-deprived conditions, a feature that is conserved in *S. aureus*. Aconitase not only negatively regulates its own expression, but also impacts the enzymes involved in both its substrate supply and product utilization. This moonlighting activity concurrently upregulates pyruvate carboxylase expression, allowing it to compensate for the TCA cycle deficiency associated with iron scarcity. These results highlight the cascade of complex posttranscriptional regulations controlling *S. aureus* central metabolism in response to iron deficiency.

## INTRODUCTION

The transition from oxidized to reduced state of iron generates a redox potential couple that is utilized by enzymes to transfer energy. Therefore, iron is an essential compound for sustaining life. Mammals use this dependence on iron in response to infections through a defense mechanism called nutritional immunity, whereby they sequester metal ions to inhibit pathogen growth (1). Bacteria adapt to iron-sequestered or -depleted host environments using sophisticated systems to capture and import iron with siderophores and dedicated transporters. However, an excess of iron is also toxic in its reduced state, which generates harmful reactive oxygen species (ROS) that can damage macromolecular compounds. Balancing the iron content is therefore crucial for organisms to optimize their fitness, as they must navigate between the need for iron and its potential toxicity. Many bacteria address this challenge by regulating the expression of iron import through the ferric uptake regulator (Fur). This protein acts as a repressor that is fully active in iron-replete cells but becomes inactivated under conditions of iron starvation (2,3). By modulating iron import, bacteria manage their iron levels to ensure proper function and minimize the detrimental effects of excess iron.

Bacterial small regulatory RNAs (sRNAs) typically bind to mRNA molecules through base-pairing interactions, thereby modulating their activity. These sRNAs play a role in maintaining general homeostasis and facilitating adaptation to fluctuating growth conditions (4). sRNAs thereby provide an additional layer of regulation that is essential for optimal fitness and adaptation to variations in iron availability. A commonly observed strategy, conserved among both Gram-positive and Gram-negative bacteria, is the production of sRNAs under iron-starved conditions. These sRNAs act by downregulating the expression of genes encoding non-essential iron-requiring enzymes, thus facilitating the conservation of iron resources for allocation to essential cellular processes (5).

*Staphylococcus aureus*, a pathogen that infects both humans and livestock, exhibits remarkable adaptability to diverse ecological niches and is responsible for numerous diseases. In a recent study, we identified IsrR, an sRNA that is repressed by Fur (6). IsrR forms base-pairing interactions with mRNAs predominantly involved in the expression of iron-containing enzymes, thereby preventing their translation. The primary function of IsrR is presumed to be conserving iron in environments lacking it, ensuring its availability for essential cellular processes.

The absence of IsrR significantly diminishes *S. aureus* virulence (6). This outcome aligns with the inability to adapt to the host iron sequestration response. Furthermore, we demonstrated that in anaerobic growth conditions, IsrR modulates nitrate respiration by targeting mRNAs associated with this metabolic pathway, which comprises 4 iron-containing enzymes. Through bioinformatics analysis, numerous targets of IsrR were predicted, indicating its potential role as a global regulator capable of rerouting metabolic pathways in response to iron starvation (6,7). Notably, some of these targets are associated with citrate metabolism.

Citrate is a key compound linking metabolic synthesis and degradation pathways, including those of carbohydrates, fatty acids, and amino acids. It is the initial molecule synthesized in the tricarboxylic acid (TCA) cycle and is formed through the condensation of oxaloacetate and acetyl-CoA by the enzyme citrate synthase. Additionally, citrate acts as a regulator for various enzymatic reactions (8). Finally, citrate is an iron chelating agent which may operate as an iron carrier (9). Given its central position in metabolism, enzymes associated with citrate are tightly regulated at multiple levels. One example of such regulation involves aconitase, a TCA enzyme that isomerizes citrate to isocitrate, with an additional moonlighting function. In several species, growth conditions characterized by iron deprivation revealed a regulatory role for aconitase, in which it exerts feedback control over its expression (10,11). In *S. aureus*, the gene *citB*, encoding aconitase, is positively regulated by the transcriptional activator CcpE, which reportedly responds to citrate levels (12-14).

In this study, we demonstrate a role for IsrR in a regulatory cascade leading to the downregulation of aconitase in *S. aureus* during iron starvation. Our findings reveal that IsrR exerts its aconitase regulatory control through both direct and indirect mechanisms. We also document the *S. aureus* aconitase moonlighting activity, showing that in iron-starved conditions, its RNA binding activity down-regulates mRNA encoding TCA-cycle enzymes. These observations lead to a detailed model for the intricate and multifaceted regulatory mechanisms that operate during iron starvation in *S. aureus*.

## MATERIAL AND METHODS

### Bacterial strains, plasmids, and growth conditions

The bacterial strains used for this study are described in Supplementary Table S1. The work was performed with *S. aureus* HG003 strain (15). Gene annotations refer to NCTC8325 nomenclature (file CP00025.1) retrieved from Genbank and Aureowiki (16). Plasmids were engineered by Gibson assembly (17) in *Escherichia coli* IM08B (18) as described (Table S2), using the indicated appropriate primers (Table S3) for PCR amplification. Plasmids were verified by DNA sequencing and transferred into HG003 or derivatives. Chromosomal mutants (point mutations, deletions and insertions) were either reported or constructed for this study (Table S1) using pIMAY derivatives as described (19). Staphylococcal strains were routinely grown in glass flasks in Brain Heart Infusion (BHI) broth (flask-to-medium ratio of 5:1) at 37°C aerobically with an agitation of 180 rpm. This last parameter is important because the effects of iron limitation can be masked by aeration limitation and affect the transcription of iron-regulated genes (20). *E. coli* strains were grown aerobically in lysogeny broth (LB) at 37°C. Antibiotics were added to media as needed: ampicillin 100 μg/ml and chloramphenicol 20 μg/ml for *E. coli*; chloramphenicol 5 μg/ml, erythromycin 1 µ/ml and kanamycin 60 μg/ml for *S. aureus*. Iron-sequestered media was obtained by adding 2,2’-dipyridyl (DIP) at a concentration of 0.5 mM and incubation for 30 min before adding bacteria. At this concentration, *S. aureus* growth is partially impacted likely due to an alteration of respiration capacity (21). At 0.5 mM DIP, the differential impact between HG003 and its isogenic *ΔisrR* derivative on growth is moderate and this is an appropriate concentration to observe *ΔisrR*-dependent phenotypes without major growth defects (Figure S1).

### Biocomputing analysis

Pairing predictions between IsrR and the mRNA targets were made using intaRNA (22) set with default parameters except for suboptimal interaction overlap set to “can overlap in both”. The sequences used for IsrR and mRNA targets were extracted from the *S. aureus* NCTC8325 strain (NCBI RefSeq NC_007795.1). For the mRNA targets, the sequences used started at the TSS when known (e.g., Exact Mapping Of Transcriptome Ends, EMOTE (23)) or were arbitrarily made to start at nucleotide -60 with respect to the +1 of translation.

### In vitro transcription, RNA labeling, and electrophoretic mobility shift assay

All RNAs were transcribed from PCR-generated DNA using MEGAscript T7 kit (Ambion). The templates for transcription were generated with forward primers containing T7 promoter sequences (Table S3) either from HG003 genomic DNA (wild-type allele) or from gBlock DNAs (Integrated DNA Technologies) containing the desired mutations. The resulting RNA sequences are indicated in Table

S4. RNAs were labeled at 5⍰.-end using [γ^32^-P]ATP (Amersham Biosciences) and T4 polynucleotide kinase (Invitrogen). Labeled and unlabeled RNAs were purified on a 5% acrylamide urea gel, eluted in elution buffer (20 mM Tris-HCl pH 7.5, 250 mM NaCl, 1 mM EDTA, 1% SDS) at 37°C and ethanol precipitated. Purified RNAs were quantified by Qubit (Thermo Fisher Scientific). Electrophoretic mobility shift assays (EMSA) were performed as described (24). RNAs were denaturated in 50 mM Tris/HEPES pH 7-7.5, 50 mM NaCl for 2 min at 80°C, followed by refolding for 10 min at 25°C after adding MgCl_2_ at final concentration of 2.5 mM. The binding reactions were in 50 mM Tris-HCl (pH 7.5), 50 mM NaCl, 2.5 mM MgCl_2_ for 20 min at 25°C. The samples were supplemented with 10% glycerol (final concentration) and loaded on a native 4% polyacrylamide gel containing 5% glycerol. The electrophoresis was performed in 0.5X Tris-borate EDTA at 4°C for 1.5 h (150 V). The results were analyzed using PhosphorImager (Amersham Biosciences).

### Western blotting

As proteins are generally stable, protein extracts were prepared by growing cells overnight. centrifugation (5 min, 4500 RPM, 4°C), and resuspension of the pellet in 400 µL of 50 mM Tris-HCl. Cells were disrupted using Fast-Prep (MP Biomedicals) and protein extracts were recovered after centrifugation (15 min, 15000 RPM, 4°C). Protein concentrations of each extract were determined by Bradford assays and 10 µg of proteins were loaded separated on a NuPAGE 4%-12% Bis-Tris gel with migration at 150 V for 45 min. Proteins were transferred from the gel to a PVDF membrane using the iBind system (Invitrogen). Immunolabeling was made overnight using a rabbit anti-Flag antibody (Invitrogen) and an anti-rabbit IgG HRP-conjugated antibody (Promega) in the iBlot system (Invitrogen). Images were obtained using a Chemidoc MP (Bio-Rad) and quantified using the ImageLab software (Bio-Rad).

### Translational reporter assay for sRNA activity

Translational reporter fusions were constructed as follows. 5’UTR regions and the first codons of the mRNA targets (14 codons for *citB* and 12 codons for *ccpE*, respectively) were fused in frame to the CDS of the fluorescent protein mAmetrine. The 5’UTRs were placed under the control of the promoter P1 from sarA (P1_sarA_). These constructs were engineered using the pRN112 plasmid and then inserted on the chromosome of *S. aureus* following the protocol from de Jong et al. (25). The insertions of the reporter fusions were confirmed by DNA sequencing.

Plasmids driving constitutive expression of *isrR* (pIsrR) and its derivatives with CRR deletions (pIsrRΔC1, pIsrRΔC2, pIsrRΔC3, pIsrRΔC1C2, pIsrRΔC1C3, pIsrRΔC2C3, and pIsrRΔC1C2C3 expressing *isrR*ΔC1, *isrR*ΔC2, *isrR*ΔC3, *isr*RΔC1C2, *isr*RΔC1C3, *isr*RΔC2C3, and *isr*RΔC1C2C3, respectively) (6,26), were introduced in strain HG003 Δ*isrR* containing the reporter fusions (Table S1). Strains harboring fluorescent reporter fusions were tested on solid or liquid media. For solid media, strains were tested on BHI plates supplemented or not with DIP and chloramphenicol when necessary. For liquid media, strains were grown overnight (2 ml of culture in 14 ml plastic Falcon test-tubes) at 37°C under agitation (180 rpm) in BHI supplemented or not with DIP and chloramphenicol when necessary. The endogenous *isrR* gene is controlled by two Fur boxes including one embedded within the transcribed sequence (6). Consequently, this Fur box was present in the series containing the *isrR*-engineered genes. To alleviate any Fur repression, experiments with pIsrR and its derivatives were therefore performed in the presence of DIP. Overnight cultures (OD_600_ 10 to 12) were then washed three times with phosphate-buffered saline (PBS 1X) and fluorescence was measured with a microplate reader (CLARIOstar), using dichroic filters with excitation wavelength 425 nm and emission wavelength 525 nm. Fluorescence was normalized by subtracting the auto-fluorescence of the HG003 strain and normalizing all cultures to 0D_600_ = 1.

### RNA preparation and transcriptome analysis

Unlike proteins, bacterial mRNAs are generally unstable. While proteins can reflect previous expression, mRNAs cannot. Moreover, the transcription of many genes is reduced or stopped during the stationary phase. For these reasons, transcriptome analysis was performed on exponentially growing cultures. *S. aureus* cultured in DIP-supplemented medium compared with non-DIP-supplemented medium reveals that over a hundred genes are differentially expressed between the two conditions (27). Most of the up-regulated genes in DIP are associated with iron or cation transport systems, while most down-regulated genes are associated with metabolic enzymes. To avoid studying a DIP effect, all transcriptomic studies (three isogenic strains) were performed simultaneously using the same DIP-supplemented medium. Overnight cultures, in biological triplicate for each strain, were diluted at OD_600_ = 0.005 in preheated BHI + DIP 0.5 mM. Cells were grown at 37°C under agitation (180 rpm) in plastic baffled flasks (flask-to-medium ratio 5:1) until OD_600_ = 1 (exponential phase). 15 mL of culture were sampled and centrifuged (10 min, 4500 rpm, 4°C), and pellets were frozen in liquid nitrogen before storage at -70°C until RNA extraction. Cell pellets were resuspended in 800 µL of lysis buffer (sodium acetate 20 mM pH 5.5, SDS 0.5%, EDTA 1 mM) and transferred into 2 mL Lysing matrix B tubes (MP Biomedicals). Bacteria were lysed using a Fastprep (MP Biomedicals) with 3 cycles of 45s at a speed of 6.5 m/s separated by incubation on ice for 1 min between each cycle. Tubes were then centrifuged (15 min, 14000 rpm, 4°C) and the RNA was extracted from the aqueous phase using phenol/chloroform as described (28). 30 µg RNA was treated using Turbo DNase I (Ambion) and purified using the RNA Clean-up and concentration kit (Norgen Biotek). RNA quality was assessed using an Agilent Bioanalyzer (Agilent Technologies) and sequenced by RNA-seq Illumina technology.

Data were processed by a Snakemake workflow (29), with FastQC (v0.11.9) quality control, creation of an index from the genome of *S. aureus* NCTC8325 Refseq NC_007795.1, mapping the reads onto the genome with Bowtie 2 (v2.4.1) (30), selection of the mapped reads with Samtools (v1.13) (31) and their counting by the FeatureCounts tool of the Subread package (v2.0.1) (32) on the coding sequences listed in genome annotation augmented with the list of sRNAs selected from a previous analysis (33), as well as differential gene expression analysis with the SARTools (v1.7.2) package (34) configured with the DESeq2 method (35). Volcano plots presenting fold-change vs significance were drawn using VolcaNoseR (36).

### Intracellular citrate quantification

Intracellular citrate concentration was measured using the Citrate Assay Kit (Abcam) following the manufacturer’s protocol. With this kit, citrate is converted to pyruvate, the latter being quantified by a probe becoming intensely colored (570 nm). Initial sample consisted of 500 µL of cells in stationary phase, and results were finally normalized at OD_600_ = 1 to account for growth differences between the samples.

### Statistical tests

All data described in this paper originates from repeated experiments. Their number is indicated in the figure legends (N = # of biological replicates). Error bars represent the standard deviation of N results. Statistical analyses between two groups were evaluated by a t-test using Excel. Relevant P-values were included in the figures using asterisk symbols (*, 0.05 ≤ p-value > 0.01; **, 0.01 ≤ p-value > 0.001; ***, p-value < 0.001).

### Quantitative reverse transcriptase PCR

Overnight cultures HG003 and its derivatives Δ*isrR, citB*(*rbp*) and Δ*isrR citB*(*rbp*) (biological triplicates, N = 3) were diluted to an OD_600_= 0.005 and incubated in BHI supplemented with DIP (0.5 mM) at 37°C. Bacterial were harvested at OD_600_ = 1. Total RNAs were extracted as described (37) and treated with DNase (Qiagen) according to manufacturer’s instructions. The last purification step was performed with the RNA Nucleospin Kit according to manufacturer’s instructions (Macherey-Nagel). The integrity of the RNAs was verified using the Agilent 2100 bioanalyzer with the RNA 6000 Nano kit (Agilent Technologies). qRT-PCR experiments and data analysis were performed as described (38). The geometric mean of the two most stable mRNAs (*ftsZ* and *gyrB* mRNAs) among eight tested were used to normalize the data. Primers used for qRT-PCRs are indicated in Table S3.

## RESULTS

### Iron sparing response regulator IsrR downregulates the amount of aconitase and CcpE

Software analysis identified several putative IsrR targets (6,7). Among these, three are associated with citrate metabolism: i) *citB* mRNA, which encodes aconitase (39), ii) *ccpE* which encodes a transcriptional activator of *citB* (12), and iii) *citM* mRNA, which encodes a putative transporter for citrate complexed with metal ions (40). These analyses also indicated that IsrR targeting of these three substrates is a conserved feature in different staphylococcal species (6,7), raising questions regarding the role of citrate in adapting to low-iron growth conditions.

The accumulation of a specific sRNA may result in changes in the stability of its target mRNAs (41). A classic approach used to identify or confirm sRNA targets involves comparing specific mRNA levels in the presence or absence of the studied sRNA (e.g. (42)). To test the activity of IsrR toward its citrate-related targets and to possibly find other IsrR targets, a transcriptomic analysis was conducted by comparing a parental strain with its Δ*isrR* derivative. Strains were cultivated in the presence of the iron chelator 2,2’-dipyridyl (DIP) to induce *isrR* expression ((6), Figure S2) and harvested in exponential growth for RNA-Seq analysis. Surprisingly, differential expression analysis using DE-seq2 (35) uncovered nearly no differentially expressed mRNAs between the parental strain and its Δ*isrR* derivative (Figure S3, Table S5). Only two genes, *mtlF* and *mhqD*, showed significant differential expression. Further bioinformatics analysis failed to identify a putative IsrR/mRNA pairing, suggesting that IsrR has an indirect effect on these mRNAs. It is worth noting that IsrR has many putative and some confirmed targets (6,26). While expression of these targets was detected, transcriptional levels remained unaffected by the absence or presence of IsrR. This observation contrasts with the typical behavior observed in Gram-negative bacteria, where the regulatory activity of an sRNA mainly influences the quantity and stability of its target mRNAs (43). The present results are supported by our previous finding that IsrR inhibits the translation of *fdhA, gltB2* and *miaB* mRNAs without significantly impacting their stability (6,26). However, we cannot exclude that under other conditions or growth phases, IsrR induction may affect target stability.

To investigate the potential effect of IsrR on the expression of *citB, ccpE* and *citM*, the three genes were engineered to encode their corresponding complete proteins extended with the 8 amino acid C-terminal Flag-tag sequence. Detection of Flag-tagged proteins using anti-Flag antibodies was used to monitor expression levels and served as a proxy for endogenous wild-type gene expression. The *citB, ccpE* and *citM* genes were replaced by their flagged alleles in strain HG003, and its isogenic derivative HG003 Δ*isrR*. The *citB-flag* and *ccpE-flag* growth curves compared to the ones on *citB* and *ccpE* defective strains (13,39) indicate that the Flag-tags did not affect their main function (Figure S4). CitM-Flag was not detected by western blot experiments, consistent with reportedly poor *citM* transcription, as tested in various conditions (7). In contrast, CitB-Flag and CcpE-Flag were detected (Figure 1). Their expression levels in HG003 cultures cultivated in a nutrient-rich medium were unaffected by the presence or absence of *isrR* as IsrR is not expressed in iron-replete media (6). We therefore added DIP to the growth medium, which alleviates Fur repression and leads to IsrR induction (6). In the presence of DIP, the abundance of CitB-Flag (Figure 1A) and CcpE-Flag (Figure 1B) is significantly lower in the parental strain compared to its Δ*isrR* derivative. These results show that the presence of IsrR correlates with a decrease of CitB and CcpE.

**Figure 1:**
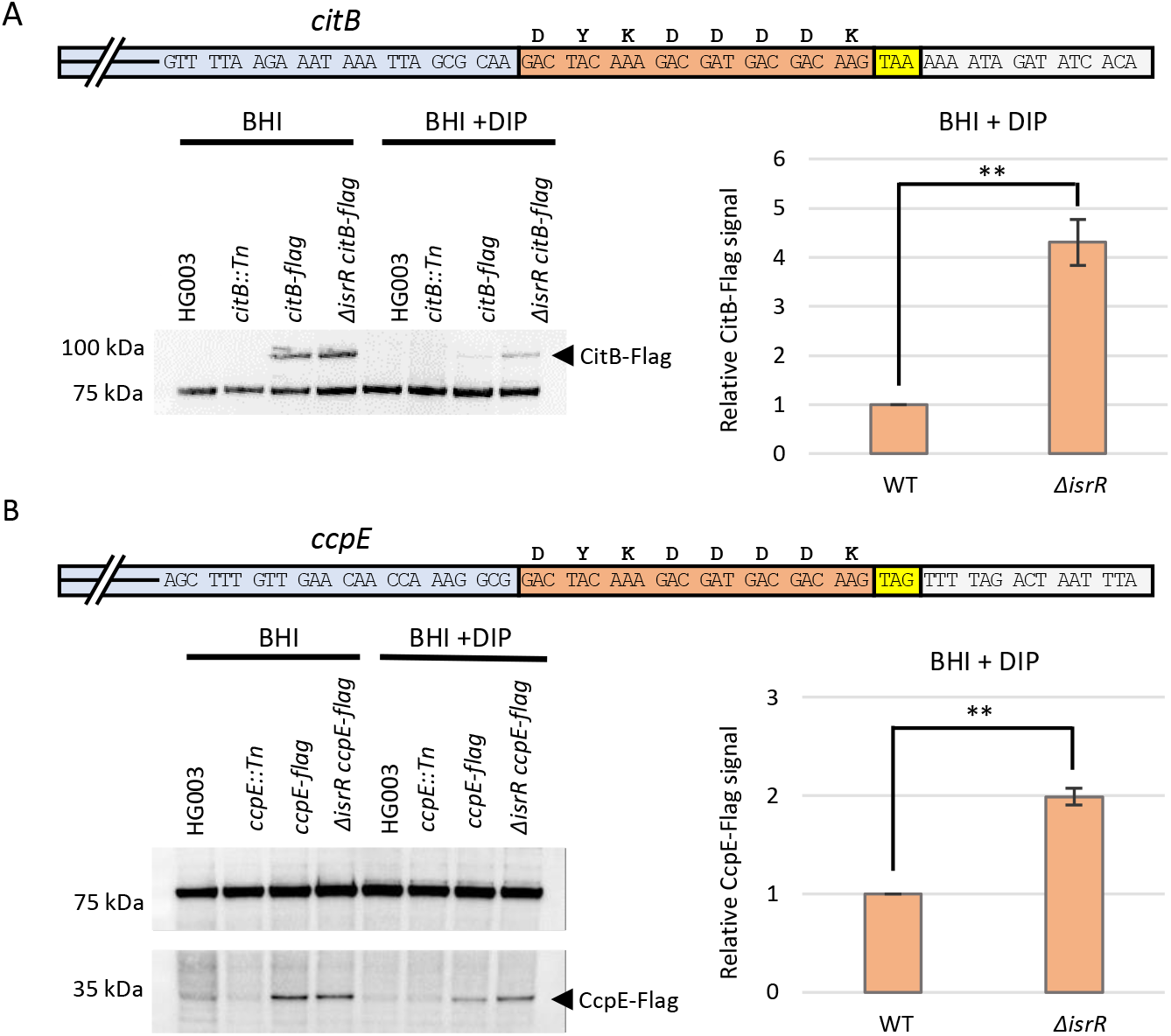
IsrR downregulates the production of CitB and CcpE. A) Upper part: schematic representation of the CitB-Flag reporter fusion. Blue, 3’ end of *citB* ORF; red, Flag sequence; yellow, native stop codon; grey, first nts of mRNA 3’UTR. Below left: western-blot experiment with anti-Flag antibodies. Genotypes and growth conditions are indicated. 100 kDa, signals corresponding to CitB-Flag; 75 kDa, non-specific signal present in all samples including Flag-less strains. The 75 kDa signal was used as a loading control for normalization. Below right: Histograms, relative quantity of CitB-FLAG in HG003 (WT) and *ΔisrR* strains grown in iron-sequestered medium (N = 3). Each specific FLAG intensity signal was normalized to the 75kDa non-specific signal and set to one for the WT strain. B) Upper part: schematic representation of the CcpE-Flag reporter fusion as in Fig. 1 A, except blue corresponds to the 3’ end of *cppE* ORF. Below left, Western-blot experiment showing the production of the CcpE-Flag as in Fig. 1A except that the CcpE-Flag band is at 35 kDa. Note a weak non-specific signal at 35kDa is also present in Flag-less strains. Below right: Histograms, relative quantity of CcpE-FLAG in HG003 (WT) and *ΔisrR* strains grown in iron-sequestered medium (N = 3). Quantification as in A).

### IsrR targets citB and ccpE 5’UTRs

Using IntaRNA software (22) with the full-length mRNA targeted sequences as input, IsrR was predicted to associate with the 5’ untranslated regions (UTRs) of *citB* and *ccpE* mRNAs, and with base-pairings occurring within the Shine-Dalgarno (SD) sequences for *citB* and *ccpE* mRNAs (Figure 2A-2B).

**Figure 2:**
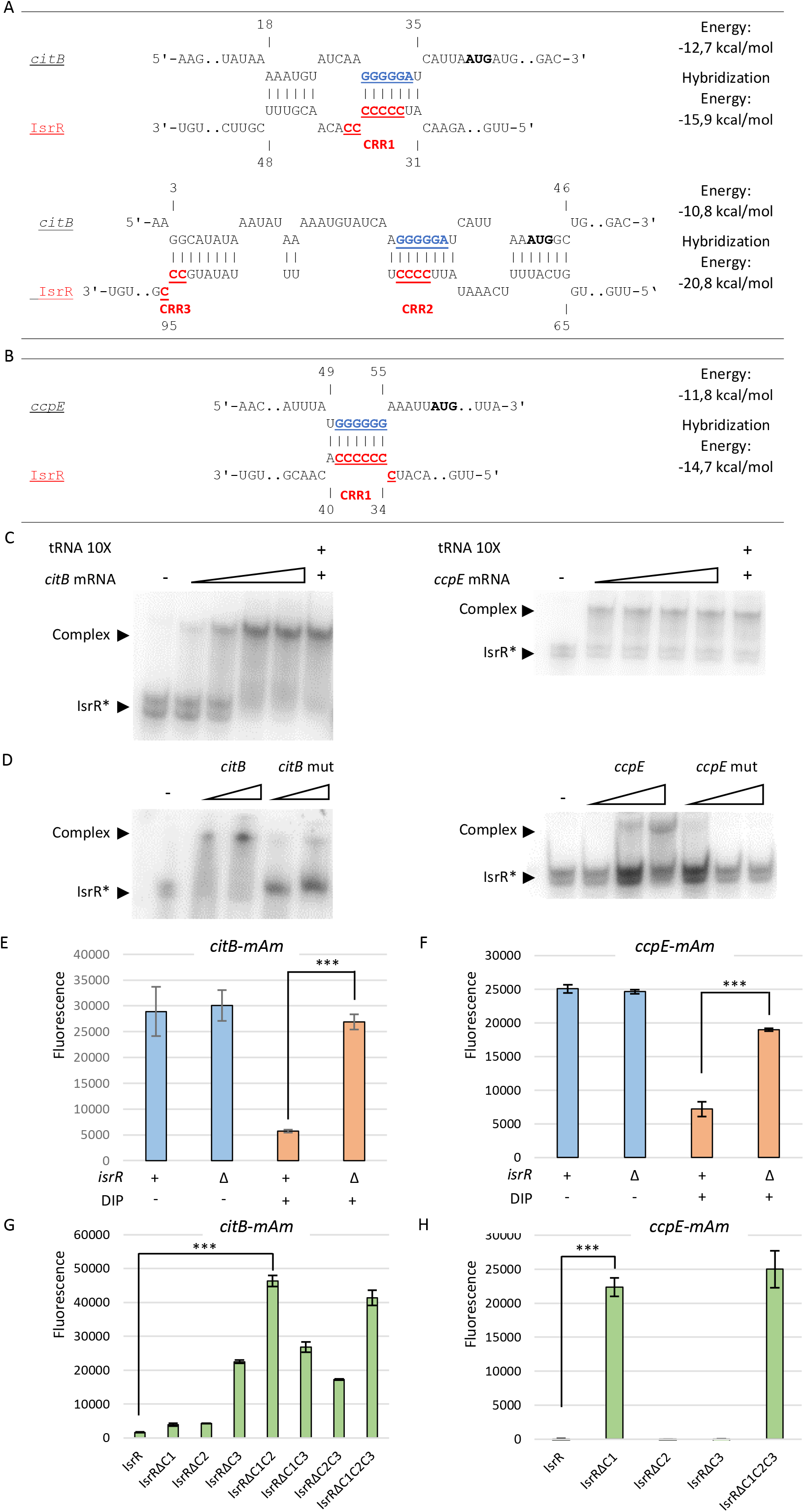
Translational repression of citB and ccpE reporters by IsrR. A) IntaRNA (18) pairing prediction between IsrR and *citB* mRNA, and B) between IsrR and *ccpE* mRNA. For A) and B) Underline blue sequences, SD; underline red sequences, CRRs; bold black characters, AUG start codon. C) Complex formation between IsrR and the *citB* and *ccpE* mRNAs. Native gel retardation assays of purified labeled IsrR with increasing amounts of either unlabeled *citB* mRNA, *ccpE* mRNA or a 10-fold excess of unlabeled synthetic RNA from *E. coli*. (D) Analysis of complex formation between IsrR with *citB* and *ccpE* mRNAs or mutated version of these mRNA targets. For sequences of RNAs generated for EMSAs, see table S4. E) *citB*-*mAm* gene expression in BHI and BHI supplemented with DIP broths. The fluorescence of overnight cultures of HG003 and HG003 *ΔisrR* derivatives harboring the fusion was determined using a microplate reader (arbitrary units). Sample fluorescence was normalized to their OD_600_ (N = 3). F) *ccpE*-*mAm* gene expression in BHI and BHI supplemented with DIP broths. Same conditions as E). G) Contribution of IsrR CRRs in *citB* regulation. The *ΔisrR* strain harboring the *citB-mAm* reporter fusion was transformed with plasmids expressing different versions of IsrR and cultured in liquid in BHI supplemented with DIP. The translational activity of the *citB-mAm* reporter fusion was quantified by measuring the fluorescence of each strain on a microplate reader (arbitrary units). Sample fluorescence was normalized to their OD_600_ (N = 3). H) Contribution of IsrR in *ccpE* regulation. The *ΔisrR* strain harboring the *ccpE-mAm* reporter fusion was transformed with plasmids expressing different versions of IsrR and cultured overnight in BHI supplemented with DIP. The translational activity of the *ccpE-mAm* reporter fusion was quantified by measuring the fluorescence (arbitrary units) of each strain on a microplate reader. Sample fluorescence was normalized to their OD_600_ (N = 3).

Electrophoretic mobility shift assays (EMSA) were performed to support bioinformatic predictions. For this purpose, we designed and produced 150 nt-long RNAs corresponding to *citB* and *ccpE* mRNAs, containing the whole predicted 5’UTRs including the ribosome binding sites (RBS) and the first 109 and 90 nts of ORFs, respectively. Duplex formation between IsrR and mRNA fragments was analysed by gel retardation assays. IsrR binds both *citB* and *ccpE* mRNA (Figure 2C); binding was specific, since a 10-fold molar excess of nonspecific synthetic RNA did not displace the *citB* mRNA from a preformed IsrR-*citB* mRNA complex. Moreover, mutations of predicted interaction sites in mRNA abolished interactions (Figure 2D), indicating that binding involves the RBSs of both mRNA targets.

We performed *in vivo* experiments to investigate whether IsrR exerts post-transcriptional control on *citB* and *ccpE* 5’UTR. To test this, the sequences corresponding to the *citB* and *ccpE* 5’UTRs, starting from their transcriptional start sites (TSS) for until the first 14 codons of *citB* and first 12 codons of *ccpE* were positioned under the control of the promoter P1_sarA_ rendering their transcription constitutive. We used a reporter fusion to evaluate *ccpE* and *citB* expression by inserting the gene encoding fluorescent protein mAmetrine (*mAm*) downstream and in frame with the beginning of each ORF (Figure S5). The reporter fusions were inserted into the chromosome at the same neutral locus (25) in both strains, HG003 and its isogenic derivative HG003 Δ*isrR*. The effects of IsrR on *citB* and *ccpE* expression were evaluated by measuring fluorescence intensity on plates and/or by quantitative determinations on microplates. To assess the effect on *ccpE* mRNA, signals were evaluated by quantitative assays, as the fluorescence from the *ccpE* 5’UTR reporter fusion was too weak to be observed on plates.

In an iron-replete medium, *isrR* being repressed by the ferric uptake regulator Fur (6), therefore, the fluorescence of each reporter was similar when introduced in both strains, HG003 and its Δ*isrR* derivative (Figures 2E-2F and S5). However, when DIP was added to the growth medium, mAmetrine fluorescence from both reporter fusions was strongly reduced in HG003 but not in HG003 Δ*isrR*. These results give strong evidence that IsrR regulates *ccpE* and *citB* post-transcriptionally by affecting their translation. They also confirm that the IsrR-dependent down-regulation of *citB* involves a direct activity of IsrR on *citB* mRNA in the tested condition.

### IsrR exerts a post-transcriptional control of *ccpE* and *citB via* its C-rich regions

A feature of many sRNAs in Gram-positive bacteria is the presence of C-rich regions (CRRs), which are likely to be seed motifs for interactions with their mRNA targets (44). *S. aureus* CRRs target G-rich regions, many of which are RBSs (45). We reported that three CRRs present in IsrR differentially contributed to the down-regulating translation of its targets (6,26). To obtain a detailed view of the regions required for IsrR activity against *citB* and *ccpE* mRNAs, we transformed the HG003 Δ*isrR* strains containing the *citB-mAm* or *ccpE-mAm* reporter fusions with plasmids expressing *isrR*, mutated *isrRs* with different CRR deletions. *isrR* and its derivatives were cloned under the control of a constitutive promoter (P_tet_). Fluorescence of the Δ*isrR* strain containing the *citB* 5’UTR reporter fusion was strongly reduced by the presence of a plasmid expressing IsrR (pIsrR) as compared to the plasmid expressing IsrR with no CRR motif (pIsrRΔC1C2C3). Deletion of CRR3 alone (pIsrRΔC3) or in combination with other CRRs (pIsrRΔC1C3, pIsrRΔC2C3) led to a moderate reduction of IsrR repression of *citB*. The reduced effect of IsrR when deleting CRR3 on *citB* may be due to lower amounts of IsrRΔC3 compared to IsrR, IsrRΔC1, and IsrRΔC2, as reported (6). Deletion of the *isrR* CRR1 (pIsrRΔC1) or CRR2 (pIsrRΔC2) did not significantly affect IsrR activity against the *citB* 5’UTR reporter, while deletion of both CRR1 and CRR2 (pIsrRΔC1C2) led to a strong fluorescence of the *citB* mAmetrine reporter (Figure 2G). Taken together, these results suggest that both CRR1 and CRR2 are independently involved in controlling *citB* mRNA translation. This conclusion is supported by IntaRNA results predicting two IsrR/*citB* mRNA pairings, one involving CRR1 and the other CRR2 and CRR3 (Figures 2A and S7). We observed a similar situation where the two CRRs of the staphylococcal sRNA RsaE could independently repress *rocF* mRNA (45). To assess which region of IsrR mediated interactions with *ccpE* mRNA, signals were evaluated by quantitative assays. Removal of CRR1 from IsrR alleviated essentially all IsrR repression of *ccpE*, while deletion of CRR2 and CRR3 had no effects (Figure 2H). Both these results and bioinformatics predictions allow us to conclude that CCR1 pairing is responsible for IsrR downregulation of *ccpE* (Figure2B).

Overall, *in vivo* and *in vitro* data indicate that IsrR binds to the Shine-Dalgarno sequences of *citB* and *ccpE* mRNAs *via* specific CRRs to prevent translation.

### Aconitase moonlighting activity down-regulates the TCA cycle, rebalancing the expression of metabolic enzymes

Aconitase catalyzes the conversion of citrate to isocitrate. However, as shown for humans and some bacteria, the apo-protein - lacking Fe-S clusters - is an RNA-binding protein (RBP) that recognizes iron-responsive elements (IREs) (46-49). We questioned if RBP activity of the apo-protein was present in staphylococcal aconitase and investigated a possible regulatory role in response to iron starvation. Essential amino acids for aconitase activities are conserved from bacteria to eukaryotes. In *Bacillus subtilis* a mutation in the aconitase ([C517A]CitB_Bs_) leads to enzymatically inactive aconitase which still binds to IREs while [R741E,Q745E]Cit_Bs_ modifications generate an aconitase defective only for its RBP activity (46,50). Aconitases from *B. subtilis* and *S. aureus* share 71% identity and their AlphaFold structures (51) are very similar, with the exception of a few minor differences at the N-terminus (Figures 3A). The region mutated in *B. subtilis* being conserved in *S. aureus* (Figure S8A), we modified the *S. aureus citB* gene to generate an aconitase deficient for its enzymatic activity, [C450S]CitB referred to as CitB(Enz), and for its RBP activity, [R734E,Q738E]CitB referred to as CitB(RBP) (Figure S8B). To maintain endogenous regulation and expression levels, the mutated alleles were introduced into the HG003 chromosomal *citB* locus by gene replacement (Table S1). In a rich medium, the enzymatically deficient *citB*(*enz*) allele was responsible for altered growth and the accumulation of intracellular citrate, similar to a strain lacking aconitase. In contrast, the *citB*(*rbp*) allele had a similar growth yield (Figure S9) and citrate amount (Figure S10) as the parental strain indicating that the strain with the *citB*(*rbp*) allele keeps its aconitase activity.

**Figure 3:**
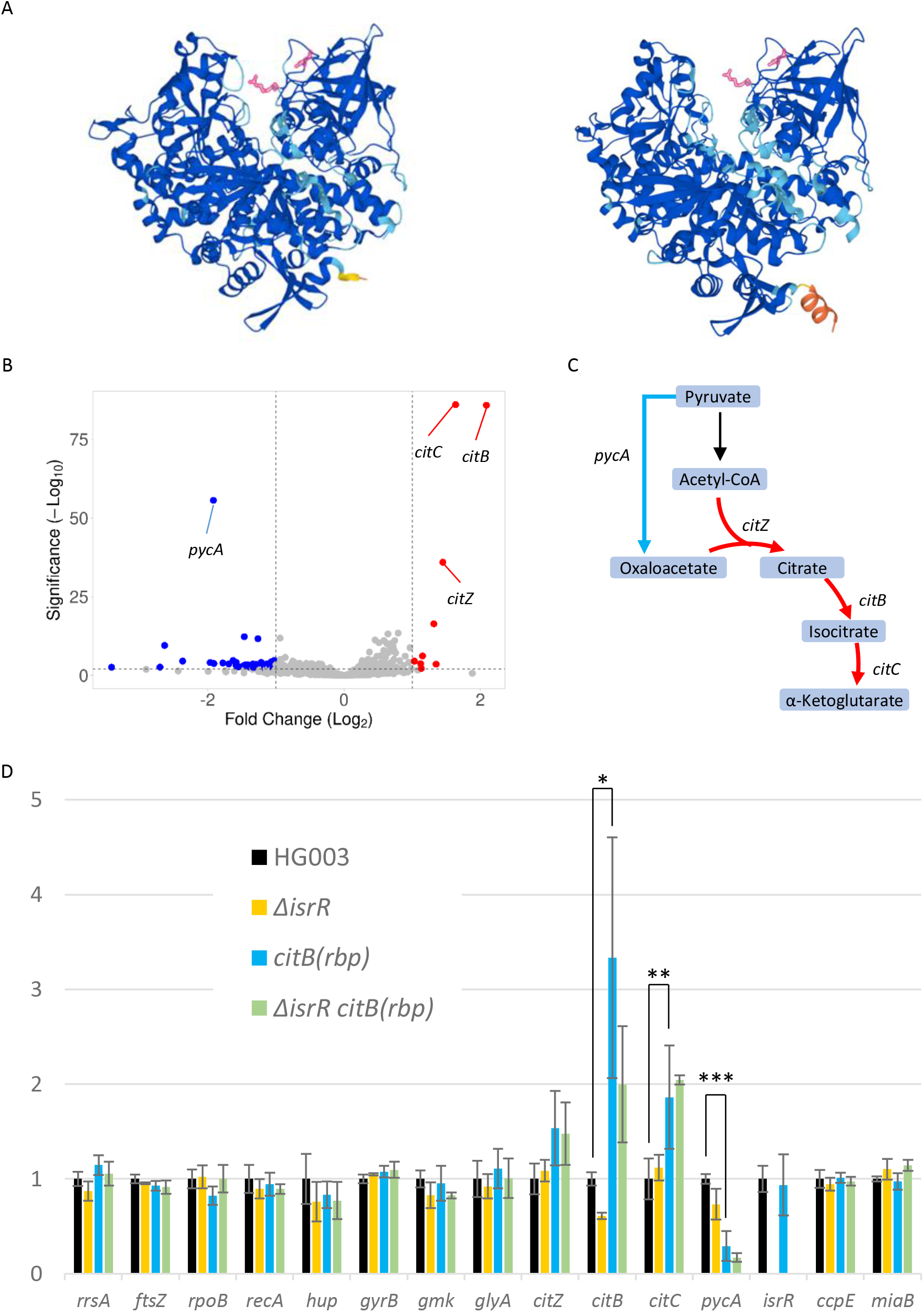
RNAs significantly affected by aconitase RBP activity. A) Alphafold structures of aconitases from *S. aureus* (left) and *B. subtilis* (right) (51). AlphaFold produces a per-residue model confidence score (pLDDT) between 0 and 100: dark blue, very high (pLDDT > 90); light blue, high (90 > pLDDT > 70); yellow, low (70 > pLDDT > 50); orange, very low (pLDDT < 50). The arginine and glutamine required for the RBP activity are shown in red. B) Volcano plot showing the relevant differences in gene expression between HG003 and *citB(rbp)* derivative upon iron starvation. The two strains were cultured overnight, followed by resuspension in fresh BHI medium supplemented with DIP. Samples were collected at an OD_600_ of 1 and RNA was extracted. Subsequently, Illumina RNA sequencing was performed, and the resulting data were analyzed using DEseq2 software. Genes with reduced expression in the *citB(rbp)* strain are shown in blue, while those with increased expression are displayed in red. Colored spots correspond to genes with a fold-change < 0.5 or >2, a significance level with a P-adjusted value below 0.005. The analysis includes the genes with a minimum of 20 reads across at least one condition (N = 3). C) Metabolic regulations mediated by apo-CitB according to transcriptomic data. Red arrows, genes downregulated by apo-CitB; blue arrow, gene activated by apo-CitB; PEP, phosphoenolpyruvate; Asp, aspartate; Lys, lysine; Asn, asparagine; Thr, threonine. D) Quantification of selected RNAs in the indicated strains relative to HG003 by qRT-PCR.

To support the putative RNA-binding deficiency of the *citB*(*rbp*) strain, and to possibly find targets of apo-aconitase, we compared the transcriptomes of HG003 with its *citB*(*rbp*) derivative. As the RBP activity is revealed only in iron-restricted growth conditions, culture media were supplemented with DIP. Experimental conditions were the same as described above (Figure 2). Transcriptomic analysis revealed that the *citB* and *citZ-citC* mRNAs were markedly upregulated in the *citB*(*rbp*) mutant (Figure 3, Table S6), thus highlighting the need for RBP activity to downregulate the expression of genes acting sequentially in the TCA cycle. The RBP-aconitase effects on gene expression shown in *S. aureus* strengthen the proposal that RBP activity is a general property of aconitases from bacteria to humans (11,46,52).

*pycA* mRNA, encoding the pyruvate carboxylase, was significantly under-represented in the *citB*(*rbp*) strain showing that this mRNA accumulates in a wild-type strain during iron-starvation (Figure 3B). PycA catalyzes the carboxylation of pyruvate to form oxaloacetate. It helps maintain the oxaloactate pool which, in addition to TCA cycle, is also used for gluconeogenesis, amino acid production and purine/pyrimidine base synthesis. Iron deprivation results in TCA cycle downregulation and, consequently, a decrease in oxaloacetate production (Figure 3C).

To support the differences observed, we compared by qRT-PCR the amounts of selected transcripts of HG003 and its *citB(rbp)* derivatives in iron-starved growth conditions (Figure 3D, Table S7). These independent experiments confirmed the accumulation of *citB, citC* and *citZ* mRNAs and the downregulation of *pycA* mRNA in the *citB(rbp)* strain when compared to its parental strain. Of note, the absence of IsrR had a moderate effect on *citB, ccpE* and *miaB* mRNAs as observed in the transcriptomic study (Figure S3 and Table S3) or reported (26).

During iron starvation, the regulatory activity of CitB due to its RBP activity plays a pivotal role in the expression levels of key genes within the citrate utilization pathway. Specifically, it facilitates the downregulation of its own corresponding mRNA, as well as that of *citZ-citC* mRNA, while concurrently upregulating *pycA* mRNA. This latter effect likely functions as a compensatory response to the deficiency in CitZ/CitB/CitC, thus ensuring the metabolic adaptation of *S. aureus* under iron-limiting conditions.

### IsrR and staphylococcal moonlighting aconitase activity can act independently to downregulate of citB expression

Could IsrR and CitB(RBP) activity against *citB* be interdependent? To address this question, the *citB(rbp)* allele was associated with the flag-tag sequence at the end of the coding sequence. The resulting allele expressed a CitB(RBP)-Flag protein, which could be monitored by Western blot experiments with anti-Flag antibodies. The endogenous citB gene was replaced by the *citB(rbp)-flag* allele in the Δ*isrR* strain and its parental strain.

In iron-sequestered growth condition, the amount of flagged aconitase was reduced when IsrR was present regardless of its allele, *citB*^+^ or *citB(rbp)*; IsrR downregulates aconitase independently of its RBP activity (Figure 4). Furthermore, CitB(RBP)-Flag was in a greater amount than CitB-Flag regardless the genetic context, *isrR*^*+*^ or Δ*isrR*. This observation shows that aconitase RBP activity downregulates aconitase independently of IsrR and also suggest that IsrR and apo-CitB act additively against *citB* mRNA.

**Figure 4:**
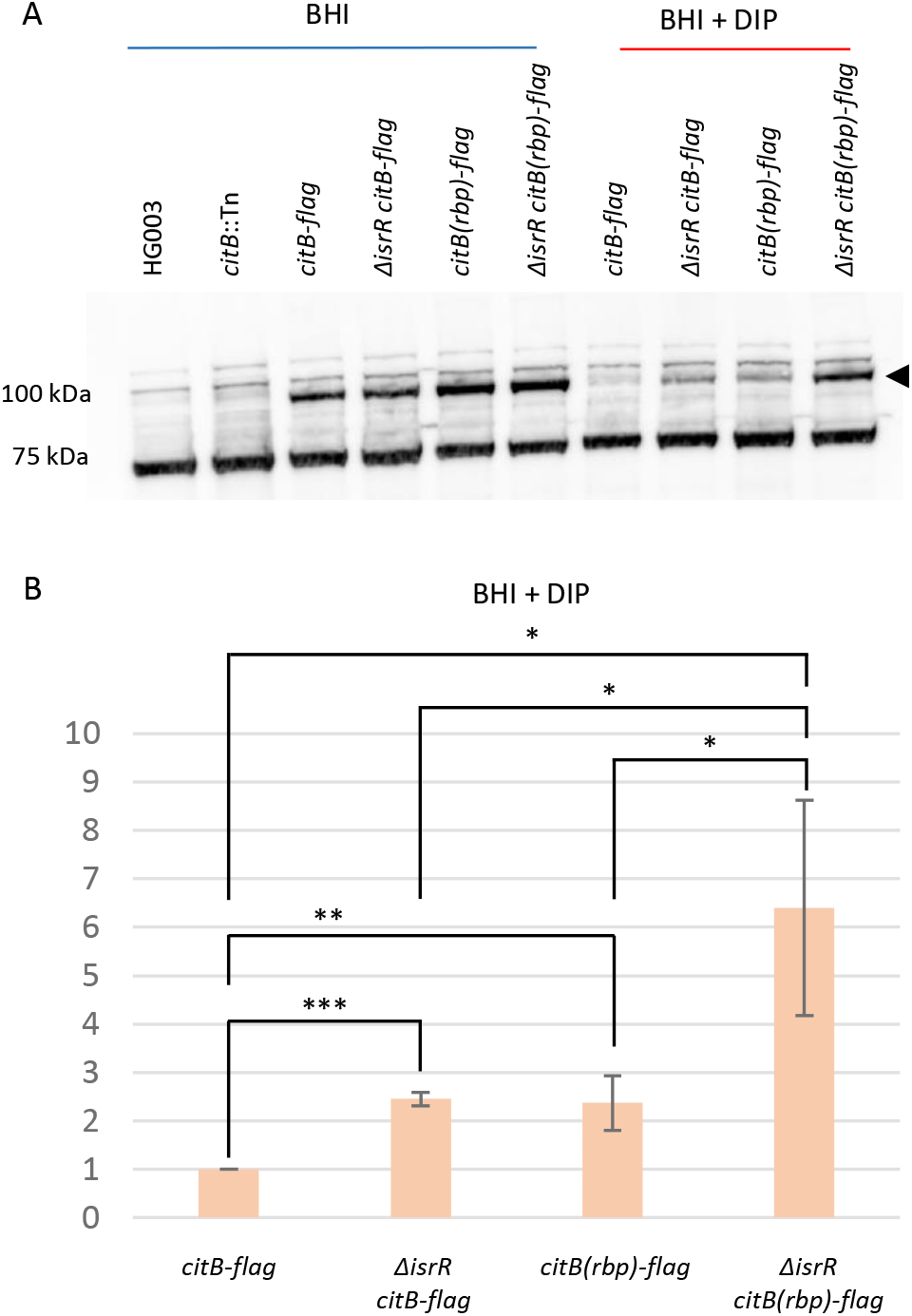
CitB RBP activity and IsrR-dependent citB regulation. A) Western-blot experiment with anti-Flag antibodies. Genotypes and growth conditions are indicated. Signals at about 100 kDa that are not present in the HG003 and HG003 *citB*::Tn strains correspond to CitB-Flag; 75 kDa, non-specific signal present in all samples including Flag-less strains. The 75 kDa non-specific signal was used as a loading control. B) Histograms of CitB-Flag and CitB(RBP)-Flag signals in BHI DIP normalized to their corresponding 75 kDa signal (N = 3).

## DISCUSSION

In response to infection, hosts activate a metal sequestration response that limits bacterial growth. We deciphered the *S. aureus* adaptive response to iron deprivation involving three post-transcriptional regulatory pathways leading to the downregulation of aconitase, which stands out as a prominent iron user (Figure 5).

**Figure 5:**
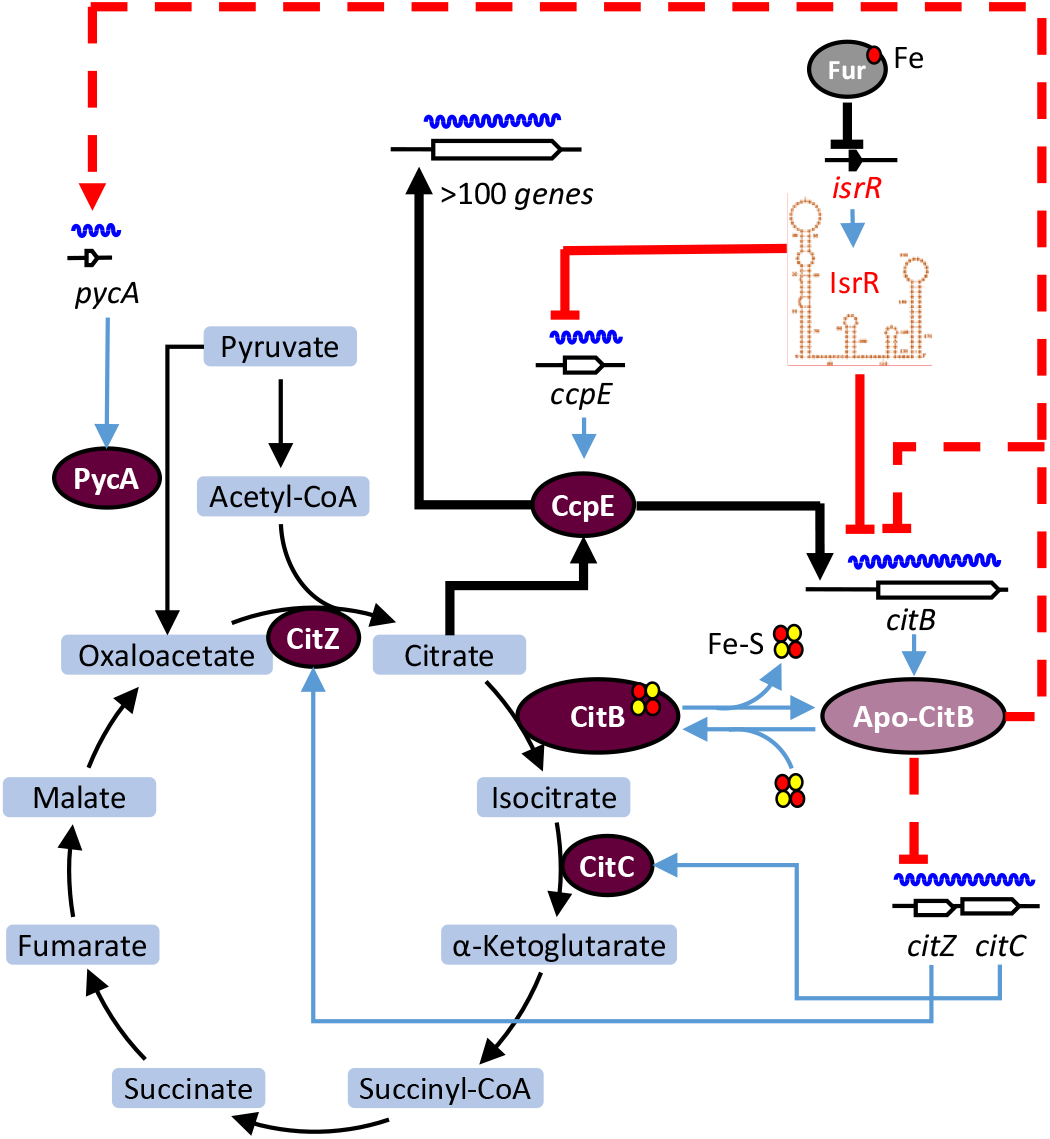
IsrR and apo-CitB controlled citrate metabolism upon iron starvation. Under iron-replete conditions, the transcription of *isrR* is repressed by Fur bound by Fe^2+^ (or Fe-S clusters (77)). Iron starvation leads to the alleviation of Fur repression, resulting in the expression of *isrR* (6). IsrR exerts its regulatory control on citrate metabolism by downregulating *citB* expression. This downregulation occurs through two distinct mechanisms: direct inhibition of *citB* mRNA translation and indirectly via inhibition of *ccpE* mRNA translation, CcpE being a transcriptional activator of *citB* and over a hundred other genes. Concomitantly, as iron being scarce, aconitase undergoes a transition from its Fe-S cluster-bound form to apo-CitB, a regulatory RNA-binding protein. Apo-CitB plays a pivotal role in the modulation of TCA cycle, leading to the downregulation of *citZ-citC* mRNA levels and its mRNA expression, while simultaneously promoting an increase in *pycA* mRNA levels, encoding pyruvate carboxylase. Red line, regulation associated with iron starvation uncovered by this study (dashed line, possible indirect regulations); blue line, transcriptions, translations or enzyme modification; thin black line, metabolic pathways; thick black line, CcpE activation or associated regulations; red disk, iron; yellow disk, sulfur.

A means of adaptation to iron-starvation conditions involves the downregulation of non-essential iron-containing proteins, which is often mediated by small non-coding RNAs induced upon iron starvation. These sRNAs comprise a diverse set, including RyhB in numerous *Enterobacteriaceae* (53), PrrFs in *Pseudomonas aeruginosa* (54), MrsI in *Mycobacterium tuberculosis* (55), IsaR1 in cyanobacterium *Synechocystis* (56), FsrA in *B. subtilis* (57) and IsrR in Staphylococcaceae (6). They play pivotal roles in coordinating the reduction of iron-utilizing pathways, with a particular emphasis on the TCA cycle (5). Aconitase is an abundant protein containing a [4Fe4S] cluster. It is no coincidence that aconitase mRNA is targeted by iron-regulated sRNAs in different species (52,53,58).

A surprising result was that the presence/absence of IsrR did not affect the quantity of its mRNA targets under the conditions tested. This result contrasts with what is generally observed on sRNA activity in Gram-negative bacteria. This difference may result from the different set of RNases or RNA chaperones they possess (59). It is noteworthy that in *B. subtilis* and *S. aureus*, the RNA chaperone Hfq lacks the overall activity observed in Gram-negative bacteria (37,60,61). Furthermore, unlike *E. coli*, transcription-translation is not coupled in *B. subtilis*, suggesting different rules for global RNA surveillance in these two species (62).

CcpE controls the expression of over 100 metabolic enzymes and virulence factors (12-14). As IsrR prevents translation of *ccpE* mRNA, transcription of CcpE-regulated genes should be affected by the presence/absence of IsrR. We did not observe significant differential expression of these genes, probably because the transcriptomic experiments were performed during exponential growth and CcpE has been reported to have major regulatory effects on the post-exponential growth phase (13).

Staphylococcal *citB* expression is activated by CcpE a citrate-responsive transcriptional regulator (14,63). Of note, CcpC, the *B. subtilis* orthologue of CcpE, has been reported to act as a repressor whose activity is relieved by citrate (64). The discrepancy between these two species was explained by the subsequent discovery that CcpC can become an activator at higher citrate concentrations (65). We show here that in *S. aureus*, iron starvation conditions promote an additional control in which IsrR downregulates CcpE, the citrate-activated aconitase transcriptional activator. The ensuing reduction of *citB* expression results in less aconitase, which should lead to accumulation of citrate (12). In turn, citrate would stimulate CcpE activity and consequently *citB* transcription, leading to a paradoxical situation where *citB* downregulation could lead to its transcriptional activation via CcpE. In iron-starved conditions, by targeting both *citB* and *ccpE* mRNAs, IsrR prevents this loophole and maintains metabolic homeostasis. The regulatory pathway comprising a three-node graph, in which A (IsrR) affects B (CcpE) and C (CitB), and B affects C, is defined as a feedforward loop (Figure S10). This organization provides additional control on the final regulation. As IsrR downregulates expression of both *citB* and *ccpE* (its transcriptional activator gene), both regulations act the same way, and the loop is defined as coherent(66). Mathematical modeling indicates that when A is a regulatory RNA (*i*.*e*., IsrR), rather than a transcriptional factor, the response to a signal (accumulation of IsrR upon iron starvation) is stronger and faster, with minimized fluctuations in target protein (*i*.*e*., aconitase) concentrations (67). These properties gained with an sRNA-controlled feedforward loop are likely critical for *S. aureus* adaptation to low iron growth conditions as encountered within the host. We show here that in *S. aureus*, iron starvation conditions promote an additional control in which IsrR downregulates CcpE, the citrate-activated aconitase transcriptional activator. The ensuing reduction of *citB* expression results in less aconitase, which should lead to accumulation of citrate (12). In turn, citrate would stimulate CcpE activity and consequently *citB* transcription, leading to a paradoxical situation where *citB* downregulation could lead to its transcriptional activation via CcpE. In iron-starved conditions, by targeting both *citB* and *ccpE* mRNAs, IsrR prevents this loophole and maintains metabolic homeostasis. The regulatory pathway comprising a three-node graph, in which A (IsrR) affects B (CcpE) and C (CitB), and B affects C, is defined as a feedforward loop (Figure S11). This organization provides additional control on the final regulation. As IsrR downregulates expression of both *citB* and *ccpE* (its transcriptional activator gene), both regulations act the same way, and the loop is defined as coherent (66). Mathematical modeling indicates that when A is a regulatory RNA (*i*.*e*., IsrR), rather than a transcriptional factor, the response to a signal (accumulation of IsrR upon iron starvation) is stronger and faster, with minimized fluctuations in target protein (*i*.*e*., aconitase) concentrations (67). These properties gained with an sRNA-controlled feedforward loop are likely critical for *S. aureus* adaptation to low iron growth conditions as encountered within the host.

A third element contributing to *citB* regulation is the aconitase protein itself, a feature conserved across diverse domains of life. Aconitase serves as an electron transfer donor through its [4Fe-4S] cluster, which are susceptible to iron deficiency and oxidative damage. Both stress conditions inactivate aconitase by disrupting these clusters. Oxidation of the [4Fe-4S] cluster to a [3Fe-4S] cluster results in the formation of an enzymatically inactive aconitase, which is prone to cluster disassembly (46,68). The fate of proteins with damage Fe-S clusters is still poorly characterized (69), as well as what happens to apo-CitB when iron is newly available. In the absence of iron, aconitase switches to function as an RBP with regulatory capabilities. In eukaryotes, the iron-responsive elements targeted by apo-CitB are hairpin-loop structures with a conserved CAGUG sequence in the loop portion (70). These sequences are not conserved in apo-CitB-regulated prokaryotic genes (10,46). However, in *E. coli*, a small hairpin-loop structure immediately downstream *acnB* mRNA was shown to be a regulatory element binding to apo-CitB (52). In *S. aureus*, mutations in the putative aconitase RBP site hinder the downregulation of *citB* mRNA, mirroring observations in *E. coli* and *B. subtilis* (46,50,52). Moreover, RBP-dependent CitB regulation extends beyond autoregulation. In eukaryotes, apo-aconitase produced upon intracellular iron starvation regulates several genes associated with iron homeostasis (71). Data on post-transcriptional control by apo-CitB in bacteria are limited. In *E. coli*, in vitro findings suggest that the translation of superoxide dismutase A (SodA) is influenced by its two apo-aconitase isoforms (72). In *B. subtilis*, besides CitB autoregulation, the production of citrate synthase (CitZ) is also inhibited by apo-aconitase (46). Citrate is not converted to isocitrate in iron-starved conditions; by preventing citrate synthase production, the regulatory activity of apo–aconitase avoids citrate accumulation. Our findings show that in *S. aureus*, this regulatory logic is conserved but with two additional features: first, *citC* mRNA, encoding isocitrate dehydrogenase (CitC), is downregulated. As this enzyme uses the product of aconitase as substrate, it should not be synthesized in conditions where aconitase is inactive. As *citZ* and *citC* are located in the same operon, they likely share the same regulatory element to achieve a coordinated regulation. In *B. subtilis*, icd (alias *citC* in *S. aureus*) is also in operon with *citZ* suggesting that both genes are also likely down-regulated by apo-CitB. Second, the TCA cycle resources biosynthetic processes, with some of its products serving as precursors for various metabolic pathways. Anaplerotic reactions are essential for replenishing depleted elements in the TCA cycle, and among these, pyruvate carboxylase (PycA) emerges as a significant enzyme that catalyzes the production of oxaloacetate (73). In addition to being a key metabolite of the TCA cycle, oxaloacetate is required for diverse biochemical pathways, including the glyoxylate cycle and amino acid synthesis. Reduced levels of CitZ, CitB, and CitC adversely impact TCA-dependent oxaloacetate production. Our findings indicate that apo-aconitase plays a role in accumulating *pycA* mRNA through a yet undiscovered mechanism. This stabilization is associated with an anticipated increase in PycA activity and subsequent oxaloacetate production. Therefore, we postulate that apo-aconitase, through its modulation of *pycA* mRNA stability, contributes to maintaining oxaloacetate pools as needed for various essential biochemical processes such as amino acid, purine/pyrimidine base synthesis and gluconeogenesis, when low iron inhibits production by the TCA cycle.

While not ortholog to IsrR, the *E. coli* RyhB regulatory RNA shares functional similarity to IsrR. *E. coli* contains two aconitases, AcnA and AcnB (74), the apoforms of both enzymes bind their cognate mRNAs (11) and RyhB downregulates the expression of both *acnA* and *acnB* (53,75). Of note, the methylcitrate dehydratase PrpD has a residual aconitase-like activity, aconitase C (76). Interestingly, there is an interplay between RyhB and the apo-aconitase RBP activity to regulate *acnB* (52). Under iron-deficient conditions, AcnB binds its mRNA to prevent RyhB-induced cleavage of its mRNA. However, our results point to a different mechanism in *S. aureus*. Firstly, IsrR activity appears to act preferentially on translation rather than on mRNA amounts and, secondly, apo-CitB does not prevent *citB* down-regulation by IsrR.

Our results demonstrate that *S. aureus* uses multifaceted mechanisms to adapt to iron-starved environments comprising a feedforward regulatory loop and the aconitase moonlighting activity, highlighting a complex interplay between post-transcriptional regulatory elements and metabolic pathways.

## ACKNOWLEDGEMENT

We thank Alexandra Gruss for her critical reading of our manuscript. We thank Patricia Kerboriou for her technical help. We thank our lab colleagues and Edel Weiss for helpful discussions and warm support. We acknowledge the sequencing and bioinformatics expertise of the I2BC High-throughput sequencing facility, supported by France Génomique (funded by the French National Program “Investissement d’Avenir” ANR-10-INBS-09). The bioinformatics analyses were performed on the Core Cluster of the Institut Français de Bioinformatique (IFB) (Funded by the “Agence Nationale pour la Recherche” ANR-11-INBS-0013).

## AUTHOR CONTRIBUTIONS

PB conceived the project. MB, SC, RHCT, CTN, EJ and PB designed experiments and interpreted the data. MB, SC and PB wrote the manuscript. MB, SC, RHCT, CTN, EJ and PB edited the paper.

## AVAILABILITY

Data generated or analyzed during this study are presented or deposited in ENA database. Strains and plasmids are available from the corresponding author on request.

## ACCESSION NUMBERS

Raw RNA-seq data have been deposited in the ENA database under accession number PRJEB74242.

