## Supplemental data for "Staphylococcal aconitase expression during iron deficiency is controlled by an sRNA-driven feedforward loop and moonlighting activity"

[1]Université Paris-Saclay, CEA, CNRS, Institute for Integrative Biology of the Cell (I2BC), 91198 Gif-sur-Yvette, France
[2]Université de Rennes 1, BRM (Bacterial regulatory RNAs and Medicine) UMR_S 1230, 35000 Rennes, France
[3]Université Paris-Saclay, Institut de Chimie des Substances Naturelles, CNRS UPR2301, 91198 Gif-sur-Yvette, France
Running Head: Aconitase post-transcriptional controls in *S. aureus*

* To whom correspondence should be addressed.

**Table S1. *Staphylococcus aureus* strains**

| Strain | Genotype | Construction/ Reference |
|---|---|---|
| **HG003** | *rsbU* and *tcaR* repaired, MSSA, Agr+ | (1) |
| **JE2** | JE2 = USA300 cured from all plasmids | (2) |
| **NE861** | USA300 *citB*::Tn | (2) |
| **NE1560** | USA300 *ccpE*::Tn | (2) |
| **SAPhB1231** | as HG003 Δ*isrR*::tag135 | (3) |
| **SAPhB2130** | as HG003 *citB*::Tn | HG003 + Φ80 lysate from NE861 |
| **SAPhB2133** | as HG003 *ccpE*::Tn | HG003 + Φ80 lysate from NE1560 |
| **SAPhB1507** | as HG003 + pP$_{isrR}$::GFP | HG003 + pP$_{isrR}$::GFP |
| **SAPhB1509** | as HG003 Δ*isrR*::tag135 + pP$_{isrR}$::GFP | SAPhB1231 + pP$_{isrR}$::GFP |
| | **Strains with P1$_{sarA}$_5'UTRcitB::*mAm* fusion** | |
| **SAPhB1755** | as HG003 P1$_{sarA}$_5'UTRcitB::*mAm* (integrated downstream of *sqr*) | HG003 + pRN112-5'citB |
| **SAPhB1759** | as HG003 P1$_{sarA}$_5'UTR*ccpe*::*mAm* (integrated downstream of *sqr*) | HG003 + pRN112-5'ccpE |
| **SAPhB1761** | as HG003 Δ*isrR*::*tag135* P1$_{sarA}$_5'UTR*citB*::*mAm* (integrated downstream of *sqr*) | SAPhB1231 +p RN112-5'citB |
| **SAPhB1765** | as HG003 *ΔisrR*::*tag135* P1$_{sarA}$_5'UTR*ccpe*::*mAm* (integrated downstream of *sqr*) | SAPhB1231 + pRN112-5'ccpE |
| **SAPhB1767** | as HG003 *ΔisrR*::*tag135* P1$_{sarA}$_5'UTR*citB*::*mAm* (integrated downstream of *sqr*) pRMC2ΔR | SAPhB1761 + pRMC2ΔR |
| **SAPhB1769** | as HG003 *ΔisrR*::*tag135 citB-mAm* pRMC2ΔR-IsrR | SAPhB1761 + pRMC2ΔR-IsrR |
| **SAPhB1771** | as HG003 *ΔisrR*::*tag135* P1$_{sarA}$_5'UTR*citB*::*mAm* (integrated downstream of *sqr*) pRMC2ΔR-IsrRΔC1 | SAPhB1761 + pRMC2ΔR-IsrRΔC1 |
| **SAPhB1773** | as HG003 *ΔisrR*::*tag135* P1$_{sarA}$_5'UTR*citB*::*mAm* (integrated downstream of *sqr*) pRMC2ΔR-IsrRΔC2 | SAPhB1761 + pRMC2ΔR-IsrRΔC2 |
| **SAPhB1775** | as HG003 *ΔisrR*::*tag135* P1$_{sarA}$_5'UTR*citB*::*mAm* (integrated downstream of *sqr*) pRMC2ΔR-IsrRΔC3 | SAPhB1761 + pRMC2ΔR-IsrRΔC3 |
| **SAPhB1900** | as HG003 *ΔisrR*::*tag135* P1$_{sarA}$_5'UTR*ccpe*::*mAm* (integrated downstream of *sqr*) pRMC2ΔR | SAPhB1765 + pRMC2ΔR |
| **SAPhB1902** | as HG003 *ΔisrR*::*tag135* P1$_{sarA}$_5'UTR*ccpe*::*mAm* (integrated downstream of *sqr*) pRMC2ΔR-IsrR | SAPhB_1765 + pRMC2ΔR-IsrR |
| **SAPhB1904** | as HG003 *ΔisrR*::*tag135* P1$_{sarA}$_5'UTR*ccpe*::*mAm* (integrated downstream of *sqr*) pRMC2ΔR-IsrRΔC1 | SAPhB1765 + pRMC2ΔR-IsrRΔC1 |
| **SAPhB1906** | as HG003 *ΔisrR*::*tag135* P1$_{sarA}$_5'UTR*ccpe*::*mAm* (integrated downstream of *sqr*) pRMC2ΔR-IsrRΔC2 | SAPhB1765 + pRMC2ΔR-IsrRΔC2 |
| **SAPhB1908** | as HG003 *ΔisrR*::*tag135* P1$_{sarA}$_5'UTR*ccpe*::*mAm* (integrated downstream of *sqr*) pRMC2ΔR-IsrRΔC3 | SAPhB_1765 + pRMC2ΔR-IsrRΔC3 |

| | | |
|---|---|---|
| **SAPhB1964** | as HG003 *ΔisrR*::*tag135* P1$_{sarA}$_5'UTR*citB*::*mAm* (integrated downstream of *sqr*) pRMC2ΔR-IsrRΔC1C2 | SAPhB1761 + pRMC2ΔR-IsrRΔC1C2 |
| **SAPhB2039** | as HG003 P1$_{sarA}$_5'UTR*citB*::*mAm* (integrated downstream of *sqr*) pRMC2ΔR-IsrRΔC1C3 | SAPhB1231 + pRMC2ΔR-IsrRΔC1C3 |
| **SAPhB2041** | as HG003 P1$_{sarA}$_5'UTR*citB*::*mAm* (integrated downstream of *sqr*) pRMC2ΔR-IsrRΔC2C3 | SAPhB1231 + pRMC2ΔR-IsrRΔC2C3 |
| **SAPhB2043** | as HG003 P1$_{sarA}$_5'UTR*citB*::*mAm* (integrated downstream of *sqr*) pRMC2ΔR-IsrRΔC1C2C3 | SAPhB1231 + pRMC2ΔR-IsrRΔC1C2C3 |
| **SAPhB2365** | as HG003 *ΔisrR*::*tag135* P1$_{sarA}$_5'UTR*citB*::*mAm* (integrated downstream of *sqr*) pRMC2ΔR-IsrRΔC1+2+3 | SAPhB1765 + pRMC2ΔR-IsrRΔC1C2C3 |
| | **Strains with *citB-flag* gene reporter** | |
| **SAPhB2054** | as HG003 *ΔisrR*::*tag135 citB-flag* | SAPhB1231 + pIMcitB-flag |
| **SAPhB2062** | as HG003 *ΔisrR*::*tag135 ccpE-flag* | SAPhB1231 + pIMccpE-flag |
| **SAPhB2075** | as HG003 *citB-flag* | HG003 + pIMcitB-flag |
| **SAPhB2077** | as HG003 *ccpE-flag* | HG003 + pIMccpE-flag |
| **SAPhB2158** | as HG003 *ΔisrR*::tag135 *citB(rbp)* | SAPhB1231 + pIMcitB(rbp) |
| **SAPhB2166** | as HG003 *citB(enz)* | SAPhB153 + pIMcitB(enz) |
| **SAPhB2167** | as HG003 *citB(rbp)* | SAPhB153 + pIMcitB(rbp) |
| **SAPhB2181** | as HG003 *ΔisrR*::tag135 *citB(rbp)-flag* | SAPhB2054 + pIMcitB(rbp)-flag |
| **SAPhB2185** | as HG003 *citB(rbp)-flag* | SAPhB2075 +pIMcitB(rbp)-flag |

**Table S2. Plasmids**

| Plasmids | Properties | Construction / reference |
|---|---|---|
| pRN112 | pJB28-NWMN29-30 + SarA_P1-mAmetrine-Term | (4) |
| pRN112-5'citB | Chromosomal integration of the *PsarA-5'citB-mAm* gene fusion between SAOUHSC_00037 and SAOUHSC_00039 | 2600/2683 on pRN112 + 2684/2685 on HG003 |
| pRN112-5'ccpE | Chromosomal integration of the *PsarA-5'ccpE-mAm* gene fusion between SAOUHSC_00037 and SAOUHSC_00039 | 2600/2683 on pRN112 + 2690/2691 on HG003 |
| pIMAY | Shuttle rep(Ts) vector in *S. aureus* | (5) |
| pIMcitB-flag | For insertion of a Flag sequence in C-ter of *citB* | 1536/1537 on pIMAY + primers 2841/2842 and 2843/2844 on HG003 |
| pIMccpE-flag | For insertion of a Flag sequence in C-ter of *ccpE* | 1536/1537 on pIMAY + primers 2849/2850 and 2851/2852 on HG003 |
| pCtl aka pRMC2ΔR | pRMC2 derivative with a *tetR* deletion for constitutive expression | (3) |
| pIsrR aka pRMC2ΔR-IsrR | pRMC2ΔR derivative. Constitutive expression of IsrR | (3) |
| pIsrRΔC1 aka pRMC2ΔR-IsrRΔC1 | pRMC2ΔR derivative. Constitutive expression of IsrRΔC1 | (3) |
| pIsrRΔC2 aka pRMC2ΔR-IsrRΔC2 | PRMC2ΔR derivative. Constitutive expression of IsrRΔC2 | (3) |
| pIsrRΔC3 aka pRMC2ΔR-IsrRΔC3 | pRMC2ΔR derivative. Constitutive expression of IsrRΔC3 | (3) |
| pIsrRΔC1ΔC2 aka pRMC2ΔR-IsrRΔC1ΔC2 | pRMC2ΔR derivative. Constitutive expression of IsrRΔC1C2 | (3) |
| pIsrRΔC1ΔC3 aka pRMC2ΔR-IsrRΔC1ΔC3 | pRMC2ΔR derivative. Constitutive expression of IsrRΔC1C3 | (6) |
| pIsrRΔC2ΔC3 aka pRMC2ΔR-IsrRΔC2ΔC3 | pRMC2ΔR derivative. Constitutive expression of IsrRΔC2C3 | (6) |
| pIsrR aka pRMC2ΔR-IsrRΔC1ΔC2ΔC3 | pRMC2ΔR derivative. Constitutive expression of IsrRΔC1C2C3 | (6) |
| pIMcitB(enz) | For insertion of the C450S mutation to inactivate enzymatic activity of CitB | 1536/1537 on pIMAY + 3010/3011 and 3012/3013 on HG003 |
| pIMcitB(rbp) | For insertion of the R724E and Q738E mutations to inactivate RNA-binding activity of CitB | 1536/1537 on pIMAY + 3006/3007 and 3008/3009 on HG003 |
| pIMcitB(rbp)-flag | For insertion of the R724E and Q738E mutations to inactivate RNA-binding activity of CitB-Flag | 3038/3039 on pIMcitB(rbp) |

| | | |
|---|---|---|
| pJET1.2 | Cloning vector: only recombinant plasmids are able to propagate | Thermo Scientific |
| pJET-T7isrR | Plasmid with template for T7 production of IsrR RNA | 2770/2771 on HG003 and ligation into pJET1.2 |
| pJET-T7isrR3Cmut | Plasmid with template for T7 production of IsrR RNA mutated in all Cs | Ligation of gBlock 3225 into pJET1.2 |
| pJET-T7isrRC1mut | Plasmid with template for T7 production of IsrR RNA mutated in C1 | Ligation of gBlock 3226 into pJET1.2 |
| pJET-T7citB | Plasmid with template for T7 production of *citB* RNA | 3221/3222 on HG003 and ligation into pJET1.2 |
| pJET-T7citBmut | Plasmid with template for T7 production of *citB* RNA with mutations complementary with IsrR3Cmut | Ligation of gBlock 3227 into pJET1.2 |
| pJET-T7ccpE | Plasmid with template for T7 production of ccpE RNA | 3223/3224 on HG003 and ligation into pJET1.2 |
| pJET-T7ccpEmut | Plasmid with template for T7 production of ccpE RNA with mutations complementary with IsrRC1mut | Ligation of gBlock 3228 into pJET1.2 |
| $pP_{isrR}$::GFP | Translational fusion between isrR promoter region and s*gfp* | (3) |

### Table S3. Primers

| Name | Sequence |
|---|---|
| 1536 | GGTACCCAGCTTTTGTTCCCTTTAGTGAGG |
| 1537 | GAGCTCCAATTCGCCCTATAGTGAGTCG |
| 1538 | TACATGTCAAGAATAAACTGCCAAAGC |
| 1539 | AATACCTGTGACGGAAGATCACTTCG |
| 2501 | AGAAAATACCGCATCAGGCG |
| 2502 | CCCCTTCTAAAGGGCAAAAGTG |
| 2529 | CAGGTCGACGGTATCGATAACT |
| 2600 | TTAGTTAATTATAACTAATTAAAAATGAGAAGTAAAC |
| 2666 | CCTTCACCTTCACCACGAACTGA |
| 2669 | ATGGAAATGAAGGCGAGGTG |
| 2670 | CACCAACACCCCATCCTAGT |
| 2673 | TTTGACAAAATGCAGGCACA |
| 2674 | TCTGGCAAGATTGTAACACCT |
| 2683 | GTTTCAAAAGGTGAAGAATTATTTACA |
| 2684 | TTCTCATTTTTAATTAGTTATAATTAACTAAAAGGCATATAAATATAAAAATGTATCAAG |
| 2685 | GTAAATAATTCTTCACCTTTTGAAACGTCAAAATGTTTTTTTGATTGCT |
| 2690 | TTCTCATTTTTAATTAGTTATAATTAACTAAAACTCATGAATGCTTGTAGC |
| 2691 | GTAAATAATTCTTCACCTTTTGAAACTAATGTTATTAGTAAACGATAGTCTTC |
| 2692 | GGAGGATGATTATTTTAGGTTTCAAAAGGTGAAGAATTATTTAC |
| 2693 | TTCACCTTTTGAAACCTAAAATAATCATCCTCCTAAGGTAC |
| 2701 | CACCACCGCATCATTTTGCACAA |
| 2705 | AACGAAACGTTGTGGGGGGG |
| 2706 | GTTATAACGTATATTGTCTTTTACGGG |
| 2752 | GGATCCACTAGTTCTAGAGCGG |
| 2753 | GGCTGCAGGAATTCGATATCAA |
| 2754 | GATATCGAATTCCTGCAGCCATCGTCTGCACCGTAATCAAGC |
| 2755 | GCTCTAGAACTAGTGGATCCGGCGACCAAATTGACAATGCAA |
| 2756 | TCGAGGTCGACGGTATCGATAA |
| 2757 | CGACTCACTATAGGGCGAATTGG |
| 2770 | TAATACGACTCACTATAGTTGAAAATGATTATCAATACCACATAG |
| 2771 | ACAAAAGCAGTAAACCTAAAGTG |
| 2830 | CCTTACGACACTTTATTTTTTACTATTTGGTACCG |
| 2831 | CCAAATAGTAAAAAATAAAGTGTCGTAAGGGTTTA |
| 2841 | CCTCACTAAAGGGAACAAAAGCTGGGTACCTGGAACGGTTGATATTGATTTA |
| 2842 | CTTGTCGTCATCGTCTTTGTAGTCTTGCGCTAATTTATTTCTTAAA |
| 2843 | GACTACAAAGACGATGACGACAAGTAAAAAATAGATATCACAGTAAAATTTT |
| 2844 | CGACTCACTATAGGGCGAATTGGAGCTCATGAAATTTCTTATTCCACAAAATA |
| 2849 | CCTCACTAAAGGGAACAAAAGCTGGGTACCCAACAACAAAAGAAAAACTAAG |
| 2850 | CTTGTCGTCATCGTCTTTGTAGTCCGCCTTTGGTTGTTCAACAAAGCTC |
| 2851 | GACTACAAAGACGATGACGACAAGTAGTTTTAGACTAATTTAAGGTTTGTAT |
| 2852 | CGACTCACTATAGGGCGAATTGGAGCTCTTGTCCAATGTGTAATAAAATAAAT |
| 2900 | CACCTTACCTGTATGATAAGTTTTGC |
| 2901 | GTAAGATATGGTGGCGAATTCCAAC |
| 2904 | GTAAATCGTACTGCCTCTGATGTG |
| 2905 | CACCTGTCCTGCTAACATCATATAGA |

| | |
|---|---|
| 2926 | CCATCGTTCAAATTTAGTTATG |
| 2927 | CATAATTCCCGTGATAAATTAC |
| 2930 | CGATAATTTAGAACAAGATTACC |
| 2931 | GAAACATATTAGCAACAAAGAG |
| 2941 | GGATCCCCTCGAGTTCATGAAAA |
| 2942 | AGTGAATTCGAGCTCAGATCTGTT |
| 2943 | AACAGATCTGAGCTCGAATTCACT |
| 2944 | TTTTCATGAACTCGAGGGGATCC |
| 2945 | TTTATATGCGCCTAAAAGAGAATATACGTTATAACAACGTT |
| 2967 | CCTGTACATAACCTTCGGGCATG |
| 2968 | GCTTCCAAGGAGCTAAAGAGGTC |
| 2969 | GTACCAGCCGTAGTTGATTTAGCTTC |
| 2970 | CCTTGCTTAGGGGCAATGATTTTAGG |
| 2971 | CGAAAACGTTACGTAAAGCGGCTG |
| 2980 | CGAAGTGATTCGTAAGGACGTCTTG |
| 2981 | ACCTTTTCCAATGACATCTGCAACAG |
| 2982 | AGACATTAGATGTCTTACAAGAAATCGTTAAAGGC |
| 3006 | CTAAAGGGAACAAAAGCTGGGTACCTCTAACCCTTATGTAATGTTAGGTGC |
| 3007 | CTGGCGCTAATTCGTTTTTAATTTCTATATTAGCAAACGTACCTCGAACC |
| 3008 | GTACGTTTGCTAATATAGAAATTAAAAACGAATTAGCGCCAGGTACTGAAGG |
| 3009 | ACTATAGGGCGAATTGGAGCTCTCAGTCACAGGTGAAATGATTCC |
| 3010 | CTAAAGGGAACAAAAGCTGGGTACCGTTTTTAAATTGGGCAACGAAAGC |
| 3011 | GATGTATTTGTTGATGATGTAATTGCTGCTATTGCAATATC |
| 3012 | GCAGCAATTACATCATCAACAAATACATCTAACCCTTATGTAATG |
| 3013 | ACTATAGGGCGAATTGGAGCTCTACCACGTCTTGAACCATATGAATT |
| 3036 | GAGTATGGAGCAACTTGCGGATTC |
| 3037 | ACCATCTAGACCAAGAGAATCAGCTG |
| 3038 | GACTACAAAGACGATGACGACAAGTAAAAAATAGATATCACAGTAAAATTTTAATCGG |
| 3039 | CTTGTCGTCATCGTCTTTGTAGTCTTGCGCTAATTTATTTCTTAAAACCA |
| 3040 | GGTTCGTTTTGACTCACTTGTTG |
| 3041 | CAAAGGTTTTGAACCGACCTATTATATC |
| 3046 | CTAAAGGGAACAAAAGCTGGGTACCTGTATATGACAATCTAGTAAAAAC |
| 3047 | CTAAAATGCCTAAAATCAAATCAAAATAACTAAGCTTTCATTAATTTTTCAAAAGCG |
| 3048 | CGCTTTTGAAAAATTAATGAAAGCTTAGTTATTTTGATTTGATTTTAGGCATTTTAGATT |
| 3049 | CACTATAGGGCGAATTGGAGCTCCATCCTATGCTAAGAAAAAGACA |
| 3050 | CGGTATGTTTAGCAACGCTGATTTAG |
| 3051 | TGGTGTTGTTGTCATTTAATCACCCATTTTCA |
| 3052 | GGTGTAGATATGACAACACCTGAA |
| 3053 | CTTAAATATCCAGATGGCGATTTCTG |
| 3062 | TGAAAATGGGTGATTAAATGACAACAACACCA |
| | **Primers for PCR products to T7-generated RNAs** |
| 3221 | TAATACGACTCACTATAGAAGGCATATAAATATAAAAATGTATC |
| 3222 | AAACTTTAGTAATACCTTGCTCTTCTAC |
| 3223 | TAATACGACTCACTATAGAACTCATGAATGCTTGTAGCC |
| 3224 | AGGTTGAGATATATATAAAATTTCAGCCGCT |
| | **gBlocks for T7-generated RNAs** |
| 3225 | TAATACGACTCACTATAGTTGAAAATGATTATCAATACCACATAGAACATCGGGGGCACAACGTTTC GTTCTTGTTGGATTGGTCATTTTCAAATATTGGGGTTTTATATGGGGGTAAAAGACAATATACGTTA TAACAACGTTTTATAAAAGCAGTAAACCCTTACGACACTTTAGGTTTACTGCTTTTGT |

| | |
|---|---|
| 3226 | TAATACGACTCACTATAGTTGAAAATGATTATCAATACCACATAGAACATCGGGGGCACAACGTTTC GTTCTTGTTGGATTGGTCATTTTCAAATATTCCCCTTTTATATGCCCGTAAAAGACAATATACGTTA TAACAACGTTTTATAAAAGCAGTAAACCCTTACGACACTTTAGGTTTACTGCTTTTGT |
| 3227 | TAATACGACTCACTATAGAAGGCATATAAATATAAAAATGTATCAACCCCCATCATTAAATGGCTGC AAATTTTAAAGAGCAATCAAAAAAACATTTTGACTTGAATGGCCAAAGTTATACTTACTATGATTTA AAAGCTGTAGAAGAGCAAGGTATTACTAAAGTTT |
| 3228 | TAATACGACTCACTATAGAACTCATGAATGCTTGTAGCCATGAAAGTTCAATAATTGAATAATTTAT CCCCCCAAATTATGAAGATTGAAGACTATCGTTTACTAATAACATTAGACGAAACGAAAACGTTACG TAAAGCGGCTGAAATTTTATATATATCTCAACCT |
| | **Primers for qRT-PCR** |
| 2227 rrsA | TGCATGGTTGTCGTCAGCTC |
| 2228 rrsA | CACCTTCCTCCGGTTTGTCA |
| 0616 ftsZ | ATCGTTATACCAAATGACCGTTTATTAG |
| 0617 ftsZ | GCGTAACACGTTGTCAGCTTCT |
| 0618 rpoB | GTGAACCATTCGATAACCGTATTTC |
| 0619 rpoB | CGCCAAGTGGTTGTTGTGTAA |
| 0622 recA | ATCGCAACCGGATCATGGT |
| 0623 recA | AGCAGCAACTGAGTCTACAACTACAATA |
| 0624 hup | CAAAACTCACTTGCTAAAGGTGAAAA |
| 0625 hup | TTGAATGCTGGAACTTTACTTGCT |
| 0628 gyrB | GCACGTGTTGCTGCGAAA |
| 0629 gyrB | CTTCAGGACTTTTACTAGAGCAATCG |
| 0632 gmk | TCCAGATGCGCTATTTATTTTCTTAG |
| 0633 gmk | GCGCTTCGTTAATACGACTTTGT |
| 0642 glyA | TGTTTGGAGCTGAACATGTCAAT |
| 0643 glyA | TGTCGCCCATTTCTAATGCA |
| 0608 citZ | ATGCGGCAAACTTCCTATATATGTT |
| 0609 citZ | CGCACAACGTGCCGTAAAT |
| 2197 citB | GCTGAAGCTGGAATGCTTGG |
| 2198 citB | TGTTGCGCCTTGTGGTAATG |
| 2207 citC | TGCTGCTAATGCTGCTCAAGA |
| 2208 citC | ATCATGCTCAGCTGGACGAGT |
| 2213 pycA | GGCTATGCCATCCAATGTCG |
| 2214 pycA | CCCGCTTGAACGATAAGCAA |
| 2217 IsrR | CCCCCACAACGTTTCGTTC |
| 2218 IsrR | AACGTATATTGTCTTTTACGGGCATA |
| 2235 ccpE | GGGTGTTCTTCTTTGATTGGACA |
| 2236 ccpE | ACCAACTTGCACTTGTATTTCAACA |
| 2241 miaB | CACAACAAGTCATCCTTGGGACT |
| 2242 miaB | CCAGATTGAACTGGCAAGTGG |

**Table S4. RNA sequences used for EMSAs**

| | |
|---|---|
| IsrR | UUGAAAAUGAUUAUCAAUACCACAUAGAACAUCCCCCCCACAACGUUUCGUUCUUGUUG GAUUGGUCAUUUUCAAAUAUUCCCCUUUUAUAUGCCCGUAAAAGACAAUAUACGUUAUA ACAACGUUUUAUAAAAGCAGUAAACCCUUACGACACUUUAGGUUUACUGCUUUUGU |
| citB | AAGGCAUAUAAAUAUAAAAAUGUAUCAAGGGGGAUCAUUAAAUGGCUGCAAAUUUUAAA GAGCAAUCAAAAAAACAUUUUGAC |
| ccpE | AACUCAUGAAUGCUUGUAGCCAUGAAAGUUCAAUAAUUGAAUAAUUUAUGGGGGGAAAU UAUGAAGAUUGAAGACUAUCGUUUACUAAUAACAUUA |
| citBmut | AA**CCG**AUAUAAAUAUAAAAAU**C**UAUCAA**CCCCC**AUCAUUAAAU**C**G**G**UGCAAAUUUUAAA GAGCAAUCAAAAAAACAUUUUGAC |
| ccpEmut | AACUCAUGAAUGCUUGUAGCCAUGAAAGUUCAAUAAUUGAAUAAUUUAU**CCCCCC**AAAU UAUGAAGAUUGAAGACUAUCGUUUACUAAUAACAUUA |

Red font, changed nucleotides.

**Table S5. HG003 *ΔisrR* vs HG003 transcriptomic analysis in iron starved growth condition**

| Id | FoldChange | padj | gene | Product |
|---|---|---|---|---|
| IsrR | 0,03 | 1,05E-89 | *isrR* | sRNA |
| SAOUHSC_02400 | 0,329 | 1,20E-04 | *mtlF* | PTS system mannitol-specific protein |
| SAOUHSC_02825 | 5 | 4,21E-03 | *mhqD* | hypothetical protein |

Padj; adjusted p-value, pink; genes overexpressed in *ΔisrR*, blue; genes underexpressed in *ΔisrR.* Only genes with fold change < 0.5 or >2, adjusted p-value < 0.005, and at least 20 reads in all conditions are shown.

**Table S6. HG003 *citB*(*rbp*) vs HG003 transcriptomic analysis in iron starved growth condition**

| Id | FoldChange | padj | gene | product |
|---|---|---|---|---|
| SAOUHSC_01801 | 3,11 | 9,6E-87 | *citC* | isocitrate dehydrogenase |
| SAOUHSC_01347 | 4,253 | 1,5E-86 | *citB* | aconitate hydratase |
| SAOUHSC_01064 | 0,265 | 2,6E-56 | *pycA* | pyruvate carboxylase |
| SAOUHSC_01802 | 2,731 | 1,2E-36 | *citZ* | citrate synthase |
| SAOUHSC_02729 | 2,491 | 4,3E-17 | | ABC transporter-like protein |
| SAOUHSC_03000 | 0,363 | 5,4E-13 | *cap1A* | capsular polysaccharide biosynthesis |
| SAOUHSC_03016 | 0,417 | 2,3E-12 | | hypothetical protein |
| SAOUHSC_00415 | 0,161 | 3,2E-10 | | transmembrane hypothetical protein |
| SAOUHSC_02025 | 2,221 | 7,1E-07 | | phi SLT orf 99-like protein |
| SAOUHSC_00560 | 0,492 | 1,9E-05 | | hypothetical protein |
| SAOUHSC_02400 | 0,324 | 2,1E-05 | *mtlF* | PTS system mannitol-specific protein |
| SAOUHSC_00962 | 0,194 | 3,1E-05 | | hypothetical protein |
| SAOUHSC_01708 | 2,046 | 3,2E-05 | *pxpA* | LamB/YcsF family protein |
| SAOUHSC_01144 | 0,477 | 5,2E-05 | *ftsL* | cell division protein |
| SAOUHSC_02992 | 0,334 | 7,8E-05 | | hypothetical protein |
| SAOUHSC_02641 | 0,43 | 8,2E-05 | *hrtB* | permease domain-containing protein |
| SAOUHSC_02832 | 0,256 | 9,1E-05 | | hypothetical protein |
| SAOUHSC_01324 | 0,475 | 9,9E-05 | | hypothetical protein |
| S414 | 0,292 | 1,2E-04 | | sRNA |
| SAOUHSC_01315 | 0,266 | 1,8E-04 | | hypothetical protein |
| SAOUHSC_02973 | 0,48 | 1,8E-04 | | hypothetical protein |
| SAOUHSC_01850 | 2,18 | 2,0E-04 | *ccpA* | catabolite control protein A |
| SAOUHSC_01944 | 0,458 | 2,1E-04 | | hypothetical protein |
| SAOUHSC_00825 | 0,31 | 2,4E-04 | | hypothetical protein |
| SAOUHSC_00465 | 0,396 | 2,5E-04 | *veg* | hypothetical protein |

| | | | | |
|---|---|---|---|---|
| SAOUHSC_00295 | 2,551 | 2,9E-04 | *nanA* | N-acetylneuraminate lyase |
| SAOUHSC_01180 | 0,409 | 3,1E-04 | | hypothetical protein |
| SAOUHSC_00556 | 0,452 | 3,1E-04 | *proP* | proline/betaine transporter |
| SAOUHSC_02993 | 0,367 | 5,0E-04 | | hypothetical protein |
| SAOUHSC_00923 | 0,376 | 5,3E-04 | *opp-3B* | hypothetical protein |
| SAOUHSC_01394 | 0,334 | 7,0E-04 | *lysC* | aspartate kinase |
| SAOUHSC_01853 | 0,471 | 7,0E-04 | | hypothetical protein |
| SAOUHSC_01142 | 0,422 | 7,0E-04 | *mraZ* | cell division protein |
| SAOUHSC_02923 | 0,361 | 8,2E-04 | *bcaP* | hypothetical protein |
| SAOUHSC_01103 | 0,383 | 8,2E-04 | *sdhC* | succinate dehydrogenase subunit |
| SAOUHSC_00843 | 0,46 | 9,5E-04 | *metP* | hypothetical protein |
| SAOUHSC_01569 | 3,209 | 1,1E-03 | | hypothetical protein |
| SAOUHSC_00064 | 0,461 | 1,4E-03 | *norG* | hypothetical protein |
| SAOUHSC_00924 | 0,355 | 1,8E-03 | *opp3C* | hypothetical protein |
| SAOUHSC_01641 | 0,342 | 2,0E-03 | *comGB* | hypothetical protein |
| SAOUHSC_02424 | 0,382 | 2,0E-03 | | hypothetical protein |
| SAOUHSC_01331 | 0,154 | 2,6E-03 | | hypothetical protein |
| SAOUHSC_00232 | 0,094 | 2,8E-03 | *lrgA* | murein hydrolase regulator |
| SAOUHSC_00347 | 0,447 | 2,9E-03 | | hypothetical protein |
| SAOUHSC_00316 | 0,398 | 3,1E-03 | *mepB* | hypothetical protein |
| SAOUHSC_00437 | 0,492 | 4,5E-03 | *treP* | hypothetical protein |
| SAOUHSC_01292 | 0,387 | 4,9E-03 | | hypothetical protein |

Padj; adjusted p-value, pink; genes overexpressed in *citB*(*rbp*), blue; genes underexpressed in *citB*(*rbp*). Only genes with fold change < 0.5 or >2, adjusted p-value < 0.005, and at least 20 reads in all conditions are shown.

**Table S7. qRT-PCR quantification of selected mRNAs in different genetic contexts.**

| | HG003 (average) | HG003 A | HG003 B | HG003 C | Δ*isrR* (average) | Δ*isrR* A | Δ*isrR* B | Δ*isrR* C |
|---|---|---|---|---|---|---|---|---|
| 2227/2228 *rrsA* | 1.000 | 0.954 | 1.089 | 0.963 | 0.871 | 0.960 | 0.901 | 0.765 |
| 0616/0617 *ftsZ* | 1.000 | 1.054 | 0.976 | 0.972 | 0.954 | 0.956 | 0.964 | 0.942 |
| 0618/0619 *rpoB* | 1.000 | 1.120 | 0.951 | 0.939 | 1.023 | 0.902 | 1.144 | 1.038 |
| 0622/0623 *recA* | 1.000 | 0.877 | 0.976 | 1.168 | 0.895 | 0.840 | 0.841 | 1.016 |
| 0624/0625 *hup* | 1.000 | 1.096 | 1.248 | 0.731 | 0.760 | 1.004 | 0.735 | 0.594 |
| 0628/0629 *gyrB* | 1.000 | 0.949 | 1.024 | 1.029 | 1.049 | 1.047 | 1.038 | 1.062 |
| 0632/0633 *gmk* | 1.000 | 0.998 | 0.915 | 1.095 | 0.828 | 0.990 | 0.726 | 0.789 |
| 0642/0643 glyA | 1.000 | 0.798 | 1.076 | 1.166 | 0.921 | 0.788 | 1.044 | 0.949 |
| 0608/0609 *citZ* | 1.000 | 1.003 | 0.848 | 1.176 | 1.085 | 0.968 | 1.109 | 1.192 |
| 2197/2198 citB | 1.000 | 0.921 | 1.048 | 1.036 | 0.610 | 0.605 | 0.579 | 0.647 |
| 2207/2208 *citC* | 1.000 | 0.896 | 0.884 | 1.262 | 1.117 | 0.976 | 1.247 | 1.146 |
| 2213/2214 *pycA* | 1.000 | 1.056 | 0.957 | 0.989 | 0.733 | 0.928 | 0.678 | 0.625 |
| 2217/2218 IsrR | 1.000 | 0.856 | 1.034 | 1.129 | 0.000 | 0.000 | 0.000 | 0.000 |
| 2235/2236 *ccpE* | 1.000 | 0.937 | 0.959 | 1.113 | 0.944 | 1.023 | 0.905 | 0.909 |
| 2241/2242 *miaB* | 1.000 | 1.030 | 0.976 | 0.994 | 1.103 | 1.224 | 1.087 | 1.008 |
| | *citB*(*rbp*) (average) | *citB*(*rbp*) A | *citB*(*rbp*) B | *citB*(*rbp*) C | *citB*(*rbp*) Δ*isrR* (average) | *citB*(*rbp*) Δ*isrR* A | *citB*(*rbp*) Δ*isrR* B | *citB*(*rbp*) Δ*isrR* C |
| 2227/2228 *rrsA* | 1,147 | 1,076 | 1,105 | 1,269 | 1,055 | 0,975 | 1,205 | 0,999 |
| 0616/0617 *ftsZ* | 0,929 | 0,930 | 0,878 | 0,981 | 0,913 | 0,993 | 0,895 | 0,855 |
| 0618/0619 *rpoB* | 0,823 | 0,872 | 0,717 | 0,891 | 1,001 | 1,128 | 1,051 | 0,846 |
| 0622/0623 *recA* | 0,943 | 0,809 | 1,007 | 1,028 | 0,894 | 0,855 | 0,877 | 0,951 |
| 0624/0625 *hup* | 0,834 | 0,954 | 0,686 | 0,886 | 0,771 | 1,009 | 0,643 | 0,707 |
| 0628/0629 *gyrB* | 1,077 | 1,076 | 1,139 | 1,019 | 1,096 | 1,007 | 1,117 | 1,170 |
| 0632/0633 *gmk* | 0,954 | 0,855 | 1,178 | 0,863 | 0,827 | 0,859 | 0,796 | 0,827 |
| 0642/0643 glyA | 1,108 | 0,880 | 1,237 | 1,249 | 1,008 | 0,809 | 1,226 | 1,033 |
| 0608/0609 *citZ* | 1,534 | 1,519 | 1,201 | 1,980 | 1,478 | 1,750 | 1,631 | 1,130 |
| 2197/2198 citB | 3,332 | 3,136 | 2,415 | 4,885 | 1,997 | 1,586 | 1,827 | 2,748 |
| 2207/2208 *citC* | 1,862 | 1,666 | 1,529 | 2,535 | 2,043 | 1,989 | 2,076 | 2,067 |
| 2213/2214 *pycA* | 0,292 | 0,322 | 0,477 | 0,162 | 0,170 | 0,166 | 0,226 | 0,132 |
| 2217/2218 IsrR | 0,936 | 0,931 | 0,672 | 1,312 | 0,000 | 0,000 | 0,000 | 0,000 |
| 2235/2236 *ccpE* | 1,012 | 0,959 | 1,069 | 1,011 | 0,975 | 0,941 | 1,030 | 0,956 |
| 2241/2242 *miaB* | 0,973 | 1,025 | 1,029 | 0,873 | 1,142 | 1,154 | 1,194 | 1,082 |

Relative variation of indicated RNAs (first column) in different strains (Δ*isrR*, *citB*(*rbp*), and *citB*(*rbp*) Δ*isrR*) compared to the parental strain HG003. The value for each biological triplicates are indicated. Fold change variation to wild-type: red > 2; blue < 0.5.

**Figure S1: Optimal DIP concentration needed for IsrR induction**

A

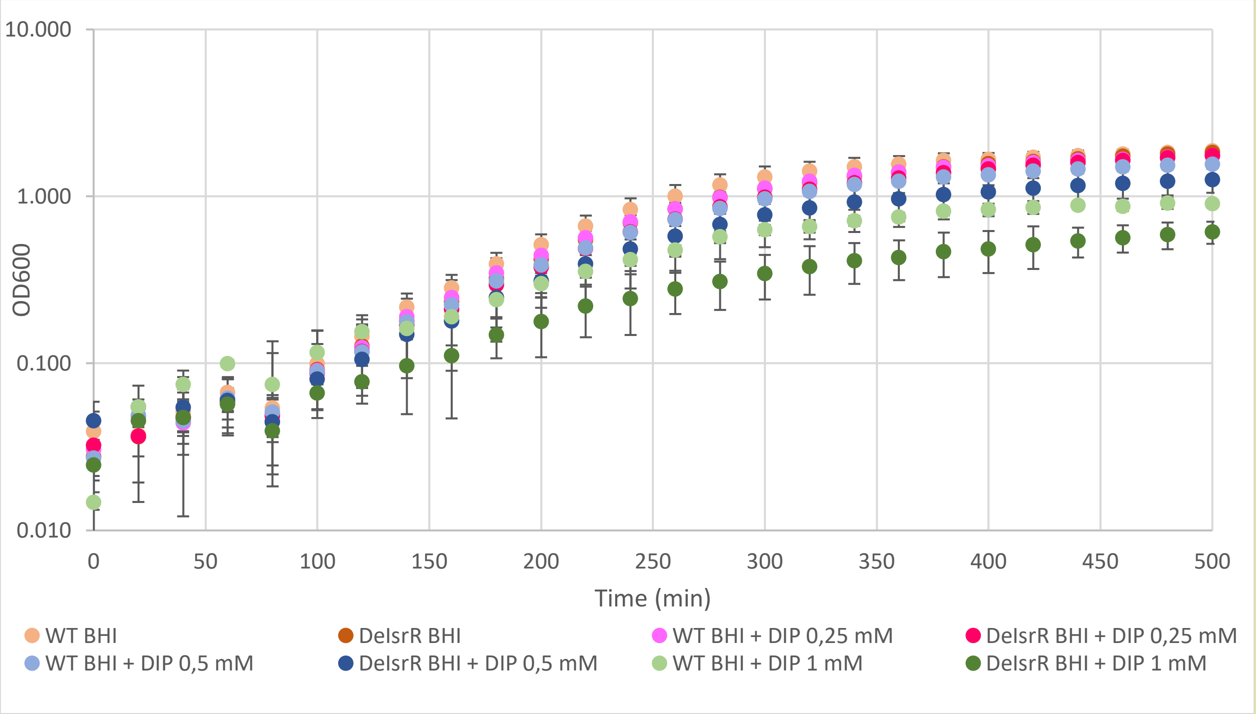

B

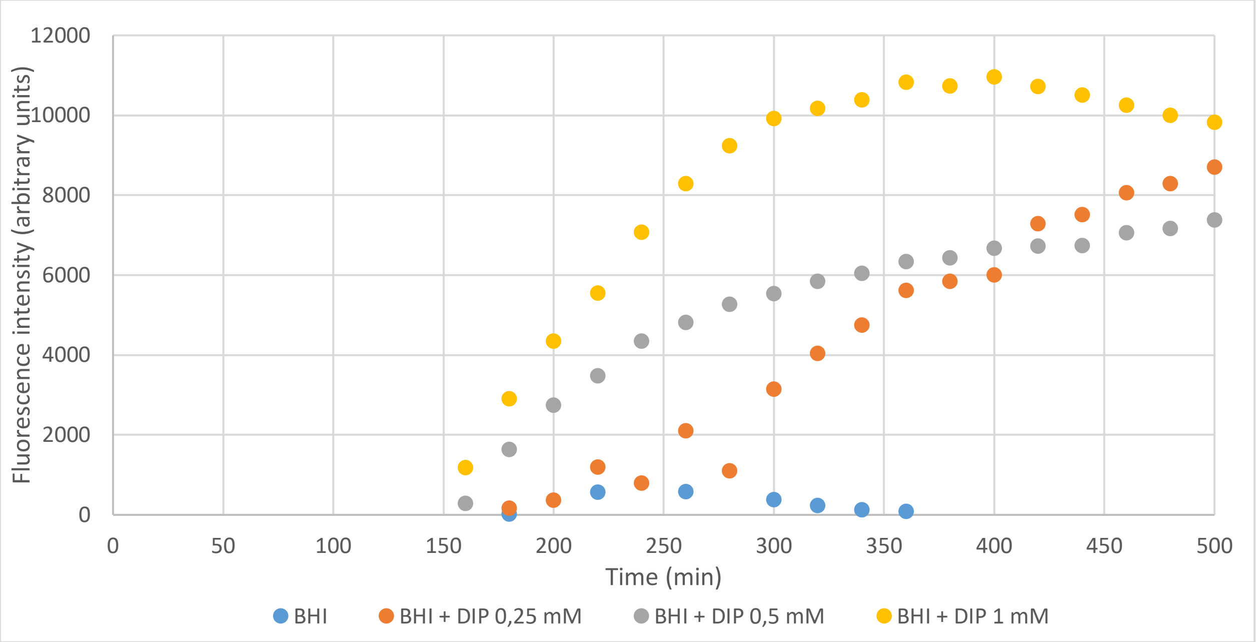

A) Growth curves of HG003 and its *ΔisrR* derivative. Strains were grown in BHI supplemented with DIP (0, 0.25, 0.5 and 1 mM) as indicated. Error bars represent the standard deviation of three independent biological replicates (N = 3). B) Fluorescence intensity (normalized to $OD_{600}$=1) of HG003 and its *ΔisrR* derivative containing the $P_{isrR}$-GFP reporter fusion (N=3) illustrating the induction of IsrR expression at the indicated DIP concentrations. OD and fluorescence intensity measurements were obtained with a Clariostar (BMG labtech) microplate reader set at 37°C with vigorous shaking (400 rpm).

**Figure S2: IsrR expression in different conditions**

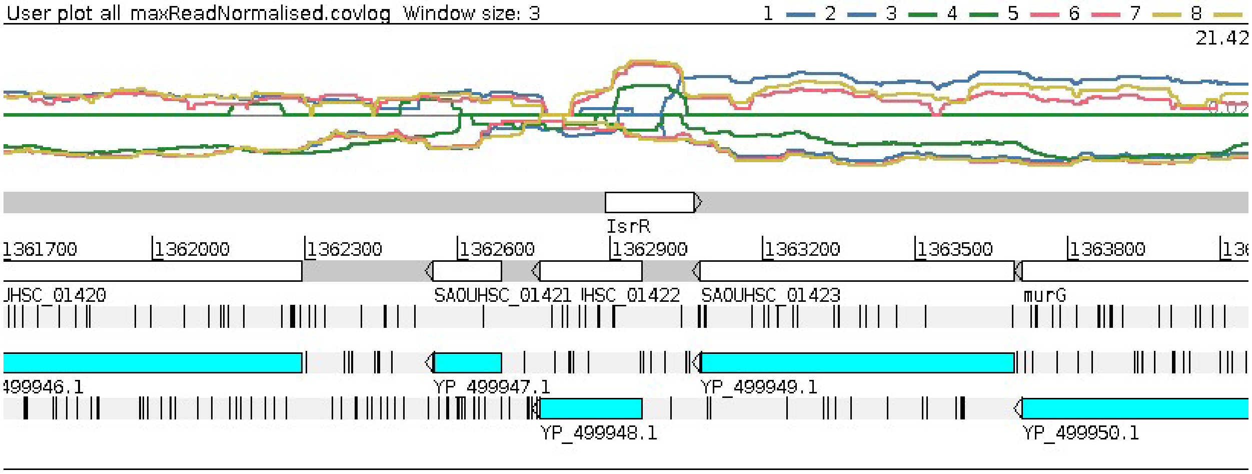

Artemis viewer (7) window showing read log-coverages from Illumina RNA-seq transcriptomes. The region covered is indicated by the genetic coordinates and gene names. Blue lines (1 and 2), Δ*isrR* strain grown in BHI DIP; green lines (3 and 4), HG003 grown in BHI; pink lines (5 and 6), HG003 in grown in BHI DIP; yellow lines (7 and 8), *citB*(*rbp*) grown in BHI DIP. Upper colored lines, + strand; lower colored lines, - strand. The normalized ratio of IsrR reads in BHI to BHI DIP is 0.01.

**Figure S3: RNAs significantly affected by the absence of IsrR**

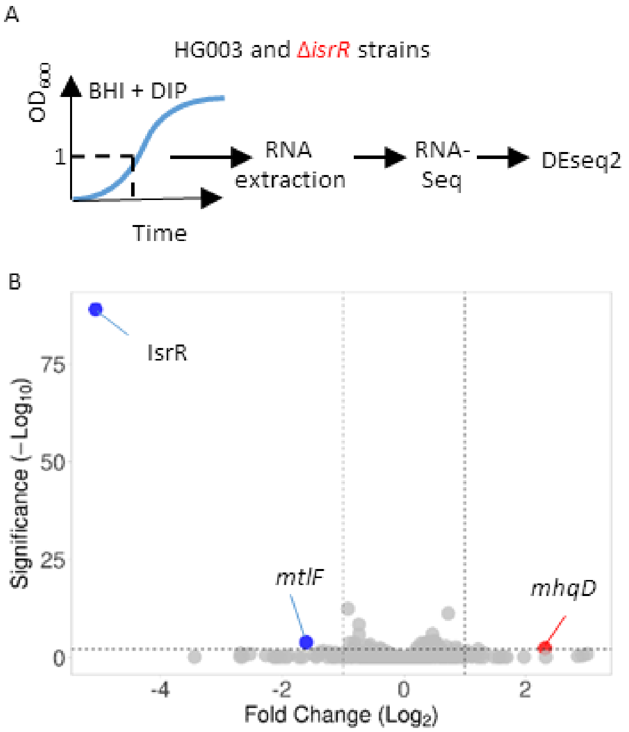

A) Experimental workflow: Schematic representation of the transcriptomic experiments. HG003 and its Δ*isrR* derivative were cultured overnight, followed by resuspension in fresh BHI medium supplemented with DIP. Samples were collected at an $OD_{600}$ of 1 and RNA was extracted. Subsequently, Illumina RNA sequencing was performed, and the resulting data were analyzed using DEseq2 software. B) Volcano plot depicting significant differences in gene expression between the HG003 strain and its Δ*isrR* derivative during iron starvation. Genes with reduced expression in the Δ*isrR* strain are shown in blue, while those with increased expression are displayed in red. Colored spots correspond to genes with a fold-change < 0.5 or >2, a significance level with a P-adjusted value below 0.005. The analysis includes the genes with a minimum of 20 reads across at least one condition (N = 3).

**Figure S4: Growth in BHI of HG003 and HG003 *ΔisrR* with *citB*⁺ *and citB-flag alleles***

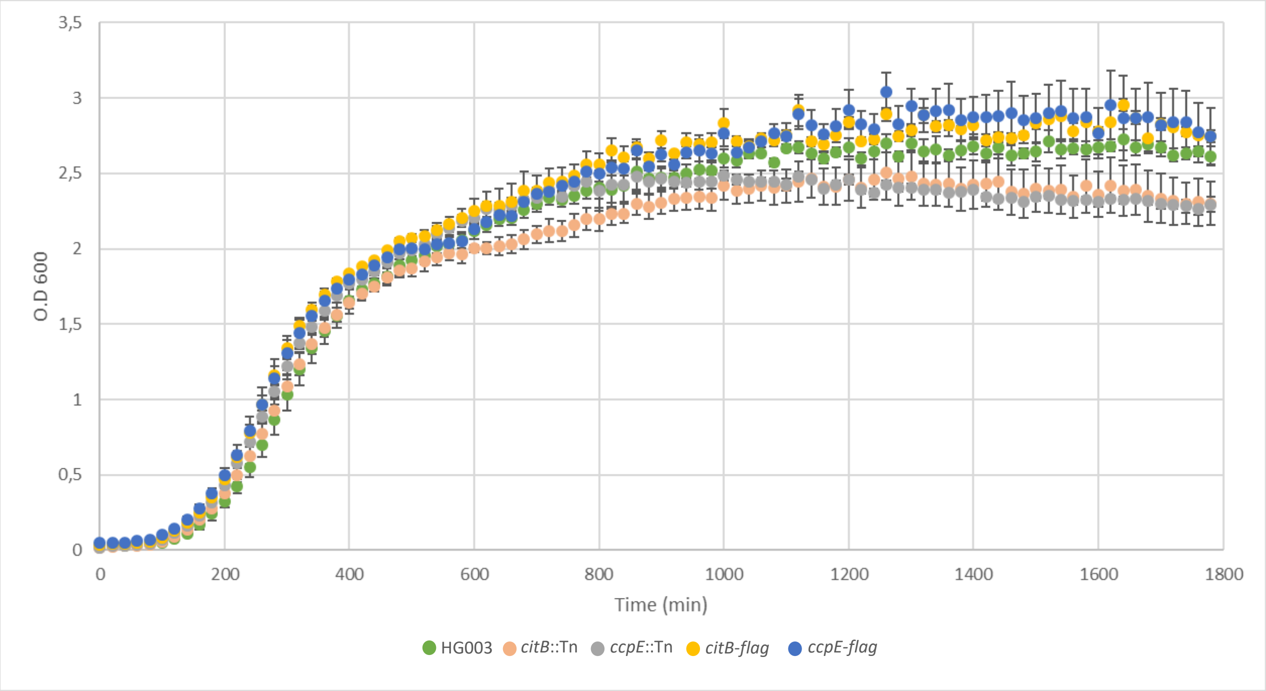

Overnight cultures were diluted 200 times and grown in rich medium (BHI) at 37°C under agitation in a microplate (Vol=200 µL) for 30 hours. Incubation and measurement of $OD_{600}$ were obtained using a Clariostar microplate reader (BMG Labtech). The HG003 *citB*::Tn and HG003 *ccpE*::Tn strains serves as a control for a non-functional CitB and CcpE. The error bars were calculated using the standard deviation of three independent biological replicates (N=3).

**Figure S5. Chromosomal reporter fusions for detection of IsrR activity**

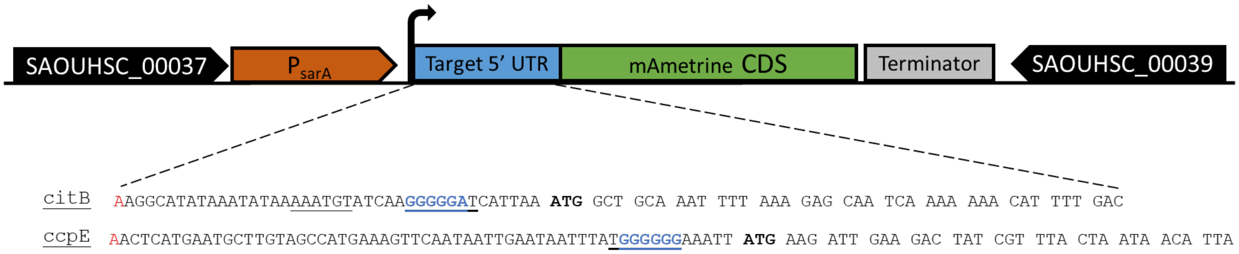

The reporter fusion consists in a region containing the 5’ UTR of each target (in blue) containing the predicted interaction region (underlined) fused in frame with the CDS of a fluorescent protein, mAmetrine (in green). This fusion is under the control of a constitutive promoter (PsarA, in orange) that allows for a strong expression of the fusion, and a strong terminator (in grey). The reporter fusions were inserted on *S. aureus* chromosome at a neutral locus between genes SAOUHSC_00037 and SAOUHSC_00039 using integrative plasmid pRN112 (4).

**Figure S6. Expression of the *citB-mAm* translational reporter in iron depleted medium**

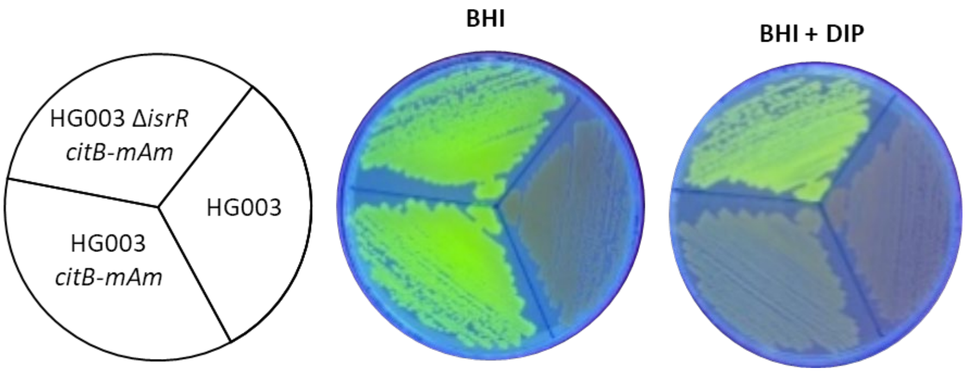

Strains were streaked out of BHI and BHI supplemented with DIP plates. Green fluorescence is visible under UV irradiation. The genotype of the corresponding strains is indicated in the cartoon on the left.

**Figure S7. IntaRNA pairing predictions**

**A**

<table>
<tr><td rowspan="2">IsrR-ΔC1</td><td rowspan="2">3 46<br>| |<br>5'-AA AAUAU AAAUGUAUCA CAUU UGCAA..GAC-3'<br>GGCAUAUA AA AGGGGGAU AAAUGGC<br>|||||||| || |||||||| |||||||<br>CCGUAUAU UU UCCCCUUA UUUACUG<br>3'-UGU..AAUGC UAAACU GUUAG..GUU-5'<br>| |<br>88 58</td><td>E= -11 kcal/mol</td></tr>
<tr><td>HE= -20.9 kcal/mol</td></tr>
<tr><td rowspan="2">IsrR-ΔC2</td><td rowspan="2">18 35<br>| |<br>5'-AAG..UAUAA AUCAA CAUUA..GAC-3'<br>AAAUGU GGGGGAU<br>|||||| |||||||<br>UUUGCA CCCCCUA<br>3'-UGU..CUUGC ACACC CAAGA..GUU-5'<br>| |<br>48 31</td><td>E= -12.8 kcal/mol</td></tr>
<tr><td>HE= -15.9 kcal/mol</td></tr>
<tr><td rowspan="2">IsrR-ΔC3</td><td rowspan="2">18 35<br>| |<br>5'-AAG..UAUAA AUCAA CAUUA..GAC-3'<br>AAAUGU GGGGGAU<br>|||||| |||||||<br>UUUGCA CCCCCUA<br>3'-UGU..CUUGC ACACC CAAGA..GUU-5'<br>| |<br>48 31</td><td>E= -12.7 kcal/mol</td></tr>
<tr><td>HE= -15.9 kcal/mol</td></tr>
</table>

**B**

<table>
<tr><td rowspan="2">IsrR-ΔC1</td><td rowspan="2">52 68<br>| |<br>5'-AAC..UAUGG AUUA UGAAG..UUA-3'<br>GGGGAA UGAAGAU<br>|||||| |||||||<br>CCCCUU ACUUUUA<br>3'-UGU..AUUUU AUAA CUGGU..GUU-5'<br>| |<br>77 61</td><td>E= -6.5 kcal/mol</td></tr>
<tr><td>HE= -13.8 kcal/mol</td></tr>
<tr><td rowspan="2">IsrR-ΔC2</td><td rowspan="2">49 55<br>| |<br>5'-AAC..AUUUA AAAUU..UUA-3'<br>UGGGGGG<br>|||||||<br>ACCCCCC<br>3'-UGU..GCAAC CUACA..GUU-5'<br>| |<br>40 34</td><td>E= -11.9 kcal/mol</td></tr>
<tr><td>HE= -14.7 kcal/mol</td></tr>
<tr><td rowspan="2">IsrR-ΔC3</td><td rowspan="2">49 55<br>| |<br>5'-AAC..AUUUA AAAUU..UUA-3'<br>UGGGGGG<br>|||||||<br>ACCCCCC<br>3'-UGU..GCAAC CUACA..GUU-5'<br>| |<br>40 34</td><td>E= -11.8 kcal/mol</td></tr>
<tr><td>HE= -14.6 kcal/mol</td></tr>
</table>

IntaRNA predictions (8) for pairings between IsrR and *citB* mRNA (A) or *ccpE* mRNA (B). Blue sequences, SD; red sequences, Cs; E, Energy; HE, Hybridization energy.

**Figure S8. Enzymatic and RBP aconitase sites**

A

```
Homo_sapiens              K-TFIYDNTEFTLAHGSVVIAAITSCTNTSNPSVMLGAGLLAKKAVDAGLNVMPYIKTSL    451
Escherichia_coli          PVDYVMNGHQYQLPDGAVVIAAITSCTNTSNPSVLMAAGLLAKKAVTLGLKRQPWVKASL    469
Staphylococcus_aureus     AEINFKDGSKATMKTGDIAIAAITSCTNTSNPYVMLGAGLVAKKAVEKGLKVPEYVKTSL    477
Bacillus_subtilis         IKFKLLNGEETVMKTGAIAIAAITSCTNTSNPYVLIGAGLVAKKAVELGLKVPNYVKTSL    471
                              . :. :  :  * :.************* *::.***:*****  **:   ::*:**

…
Homo_sapiens              RRGNDAVMARGTFANIRLLNRFLNK-QAPQTIHLPSGEILDVFDAAERYQQAGLPLIVLA    750
Escherichia_coli          RRGNHEVMMRGTFANIRIRNEMVPGVEGGMTRHLPDSDVVSIYDAAMRYKQEQTPLAVIA    768
Staphylococcus_aureus     RRGNHEVMVRGTFANIRIKNQLAPGTEGGFTTYWPTNEVMPIFDAAMKYKEDGTGLVVLA    777
Bacillus_subtilis         RRGNHEVMMRGTFANIRIKNQIAPGTEGGFTTYWPTGEVTSIYDACMKYKEDKTGLVVLA    771
                          ****. ** ********: *.:    :.  * : * .::  ::**. :*::    * *:*
```

B

| | | | | | | | | | |
|---|---|---|---|---|---|---|---|---|---|
| CitB | A | I | T | S | C | T | N | T | S |
| *citB* | GCA | ATT | ACA | TCA | TGT | ACA | AAT | ACA | TCT |
| CitB(ENZ) | A | I | T | S | **S** | T | N | T | S |
| | GCA | ATT | ACA | TCA | TCA | ACA | AAT | ACA | TCT |

| | | | | | | | | | |
|---|---|---|---|---|---|---|---|---|---|
| CitB | N | I | R | I | K | N | Q | L | A |
| *citB* | AAT | ATA | CGT | ATT | AAA | AAC | GAA | TTA | GCG |
| CitB(RBP) | N | I | **E** | I | K | N | **E** | L | A |
| *citB*(*rbp*) | AAT | ATA | GAA | ATT | AAA | AAC | GAA | TTA | GCG |

A) Clustal alignment of aconitases sequences from *H. sapiens*, *E. coli*, *S. aureus* and *B.T subtilis* (selected regions). Red, amino acids that were changed. B) Protein and DNA sequences of CitB/*citB* and mutated CitB/*citB* (selected region). Upper panel CitB(ENZ)/*citB(enz)*, lower panel CitB(RBP)/*citB*(*rbp*).

**Figure S9. Growth in BHI of HG003 and *citB* mutant derivatives**

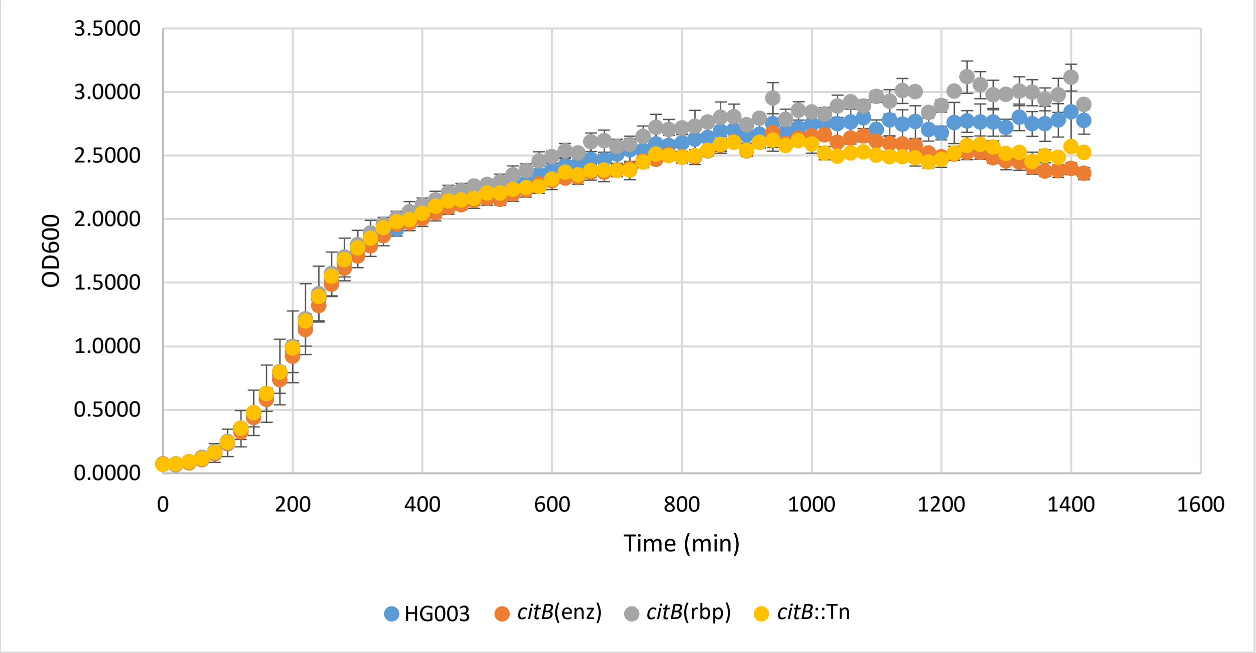

Overnight cultures were diluted 100 times and grown in rich medium (BHI) at 37°C under agitation in a microplate (Vol=200 µL) for 24 hours. Incubation and measurement of OD600 were obtained using a Clariostar microplate reader. The HG0003 *citB*::Tn strain serves as a control for a non-functional CitB. The error bars were calculated using the standard deviation of three independent biological replicates (N = 3).

**Figure S10. Intracellular citrate concentration in HG003 and *citB* mutant derivatives**

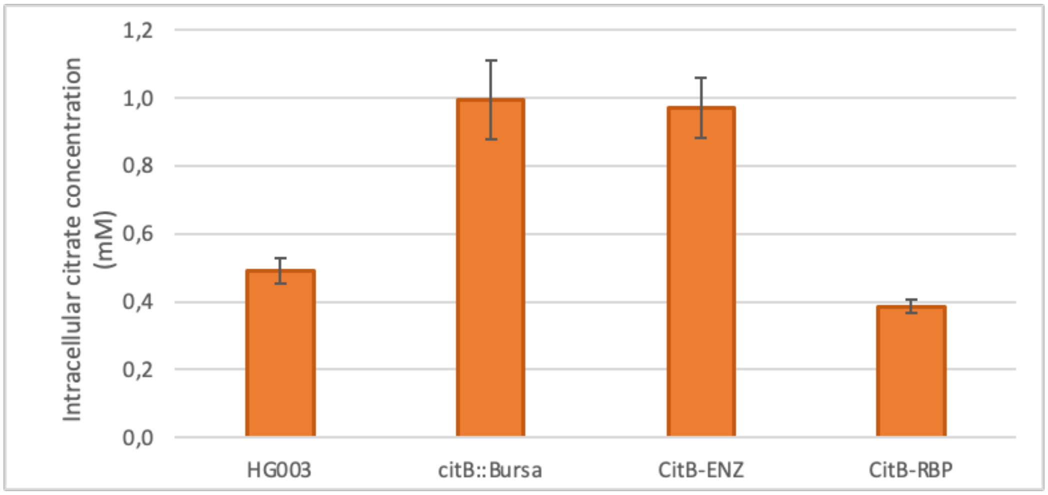

Error bars represent the standard deviation of three independent biological replicates (N = 3). The intracellular citrate concentration was obtained from 500 µL of cells in stationary phase normalized to $OD_{600}$=1 to account for the differences in cell density between the samples.

**Figure S11: IsrR feedforward loop driving *citB* regulation**

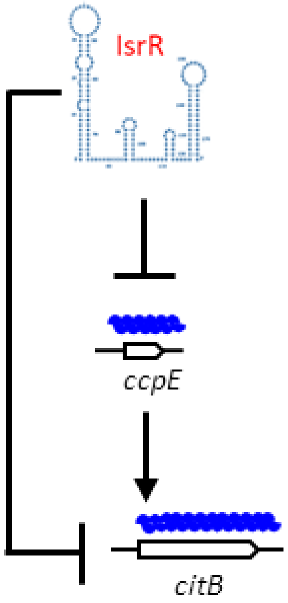

Coherent sRNA-mediated feedforward loop (smFFL) type 2 as established (9,10).
